# Dissecting dynamics and differences of selective pressures in the evolution of human pigmentation

**DOI:** 10.1101/253963

**Authors:** Xin Huang, Yungang He, Sijia Wang, Li Jin

## Abstract

Human pigmentation is a highly diverse and complex trait among populations, and has drawn particular attention from both academic and non-academic investigators for thousands of years. Previous studies detected selection signals in several human pigmentation genes, but few studies have integrated contribution from multiple genes to the evolution of human pigmentation. Moreover, none has quantified selective pressures on human pigmentation over epochs and between populations. Here, we dissect dynamics and differences of selective pressures during different periods and between distinct populations with new approaches. We propose a new model with multiple populations to estimate historical selective pressures by summarizing selective pressures on multiple genes. We use genotype data of 19 genes associated with human pigmentation from 17 datasets, and obtain data for 2346 individuals of six representative population groups from worldwide. Our results quantify selective pressures on light pigmentation not only in modern Europeans (0.0249/generation) but also in proto-Eurasians (0.00665/generation). Our results also support several derived alleles associated with human dark pigmentation may under directional selection by quantifying differences of selective pressures between populations. Our study provides a first attempt to quantitatively investigate the dynamics of selective pressures during different time periods in the evolution of human pigmentation, and may facilitate studies of the evolution of other complex traits.

**Author Summary:** The color variation of human skin, hair, and eye is affected by multiple genes with different roles. This diversity may be shaped by natural selection and adapted for ultraviolet radiation in different environments around the world. As human populations migrated out from Africa, the ultraviolet radiation in the environment they encountered also changed. It is possible that the selective pressures on human pigmentation varied throughout human evolutionary history. In this study, we develop a new approach and estimate historical selective pressures on light pigmentation not only in modern Europeans but also in proto-Eurasians. To our best knowledge, this is the first study that quantifies selective pressures during different time periods in the evolution of human pigmentation. Besides, we provide statistical evidence to support several genes associated with human dark pigmentation may be favored by natural selection. Thus, natural selection may not only affect light pigmentation in Eurasians, but also influence dark pigmentation in Africans.

## Introduction

Human pigmentation—the color of human skin, hair, and eye—is one of the most diverse traits among populations. Its obvious diversity has attracted attention from both academic and non-academic investigators for thousands of years, as noted by Charles Darwin one century ago [1] and as noticed by ancient Egyptians more than 4000 years ago [2]. Why human pigmentation diverges, however, remains a central puzzle in human biology [3]. Some researchers have proposed that the diversity of human pigmentation is adapted for ultraviolet radiation (UVR) and driven by natural selection [4–6]. Dark skin may prevent sunburn amongst individuals in low latitude areas with high UVR, while light skin may protect infants against rickets in high latitude areas with low UVR [5–9]. A better understanding of how natural selection shapes the diversity of human pigmentation could provide relevant and beneficial information for public health [4–6].

During the last decade, many studies have applied statistical tests to detect signals of natural selection in several human pigmentation genes [10–18]. These genes encode different proteins, including: signal regulators—ASIP, KITLG, MC1R—stimulating the melanogenic pathway; possible enhancers—BNC2, HERC2—regulating pigmentation gene expression; important enzymes—TYR, TYRP1, DCT—converting tyrosine into melanin; putative exchangers—OCA2, SLC24A4, SLC24A5, SLC45A2, TPCN2—controlling the environment within melanosomes; and an exocyst complex unit and molecular motor—EXOC2, MYO5A—conveying vesicles and organelles within the cytoplasm [19–33]. These proteins work at different stages of the melanogenic pathway, illustrating that human pigmentation is a complex trait affected by multiple genes with different roles.

Previous studies applied two groups of methods to detect natural selection. One group of methods detects unusually long extended haplotype homozygosity [10–16, 34, 35]. The other group of methods identifies extremely local population differentiation [10, 12–15, 17]. By applying both groups of methods, previous studies have aimed to interpret the evolution of individual pigmentation genes; however, few studies have integrated contributions from multiple genes to the evolution of human pigmentation. Moreover, none of these studies have quantitatively investigated the historical selective pressures of pigmentation genes during different epochs, and compared the differences of selective pressures between different populations. Therefore, it is necessary to perform an extensive quantification of selective pressures on human pigmentation using a creative approach.

## Results and Discussion

### The model

On the basis of a previous study [36], we measure selective pressures by (genic) selection coefficients. For any single nucleotide polymorphism (SNP) *L*, we can estimate the expectation of the selection (coefficient) difference per generation between populations *i* and *j* by

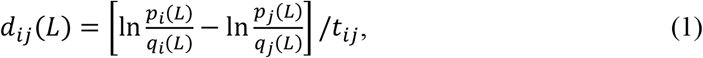

where *p* and *q* are the frequencies of derived and ancestral alleles in a population, respectively; and *t_ij_* is the divergence time of the populations *i* and *j* from their most recent common ancestor. Details of the calculations are described elsewhere [36].

We can extend Eq. 1 to summarize selection differences of multiple SNPs, because a complex trait is usually affected by multiple loci. Here, we take two bi-allelic SNPs as an example. We can estimate the selection difference of a trait affected by two SNPs between populations *i* and *j* using

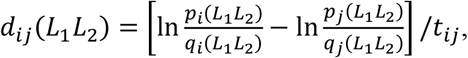

where *p*(*L*_1_*L*_2_) is the frequency of the combination carrying two alleles associated with one possible outcome of a trait, such as light pigmentation; *q*(*L*_1_*L*_2_) is the frequency of the combination carrying two alleles associated with another outcome of the same trait, such as dark pigmentation. With linkage equilibrium between *L*_1_ and *L*_2_, we have *p*(*L*_1_*L*_2_) = *p*(*L*_1_)*p*(*L*_2_) and *q*(*L*_1_*L*_2_ = *q*(*L*_1_)*q*(*L*_2_). Thus,

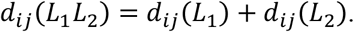

We further assume the distribution of selection difference in each SNP is identical and independent. Therefore, the variance of *d_ij_*(*L*_1_*L*_2_) is

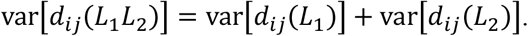

The confidence interval (CI) of *d_ij_*(*L*_1_*L*_2_) becomes 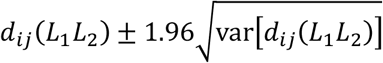.

More generally, for a trait with *n* SNPs in linkage equilibrium as well as independent and identically distributed selection differences, the expectation and variance of the total selection difference in this trait is

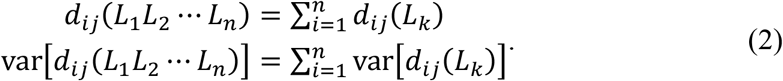

Based on Eq. 2, we develop a new approach for dissecting historical selective pressures over epochs of the human evolutionary history. We simplify the evolutionary history of six representative human populations as a binary tree (Fig. 1).

**Fig. 1.**
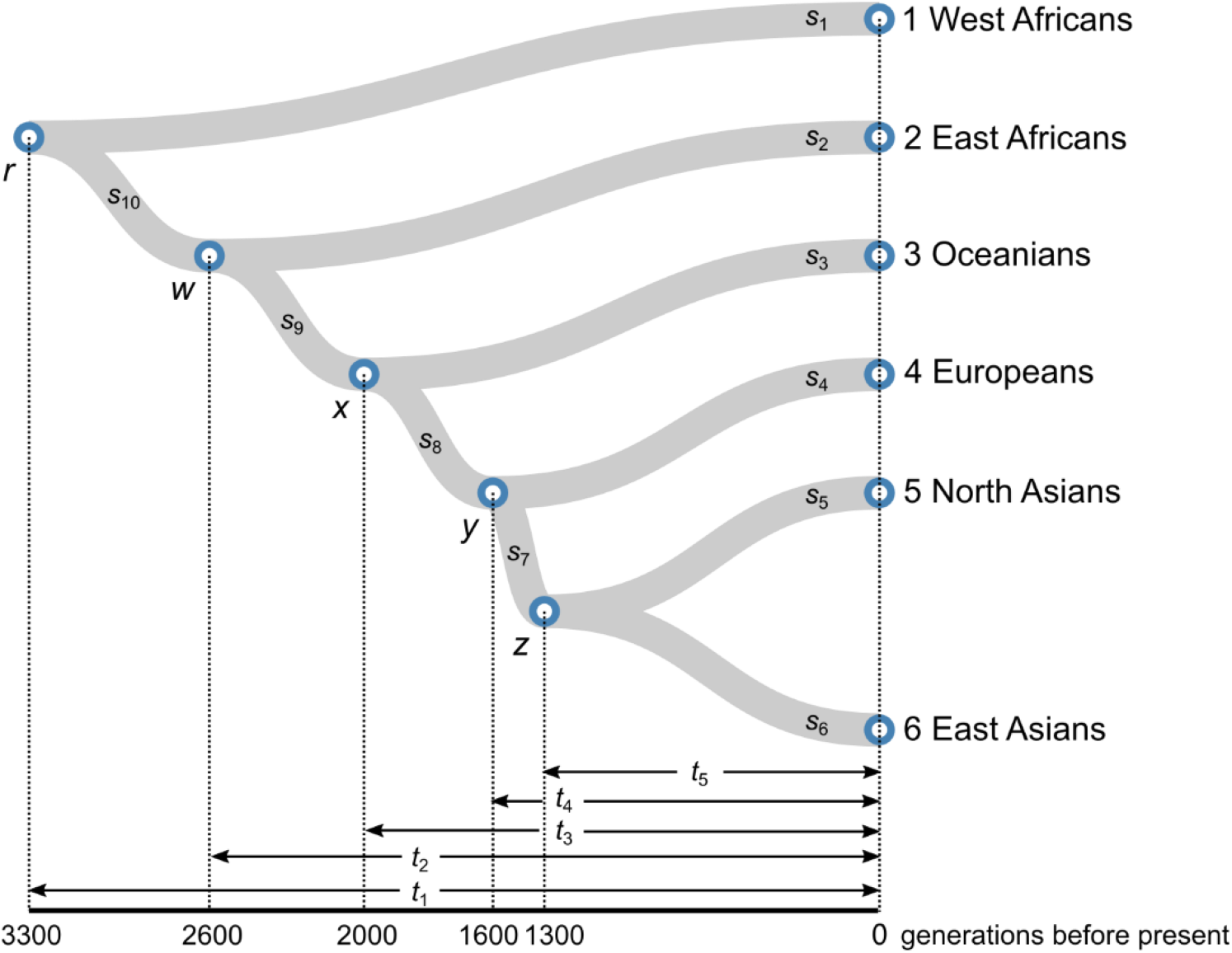
Time-varied selective pressures on an evolutionary tree. We model the evolutionary history of six representative human populations as a binary tree. Here, *s_i_* (*i* = 0, 1,…, 10) denotes the selection coefficient of the *i*-th epoch and can be obtained by estimating selection differences between paired populations. The numbers on the branches indicate different epochs. Divergence times between populations were based on previous studies [37–40]. We assumed one generation time is ~30 years. We obtain an optimal solution deviated least from neutral evolution using a probabilistic approach. The optimal solution (/generation) is 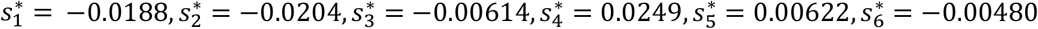, 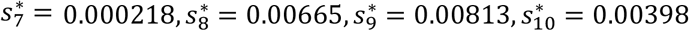. In the solution, numbers larger than zero suggest natural selection favored light pigmentation, while those less than zero indicate natural selection preferred dark pigmentation.

When *k* is the most recent common ancestor of *i* and *j*, we can divide *d_ij_* in Eq. 1 into separate terms:

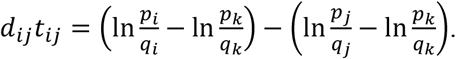

We can further divide the selection difference between paired populations into multiple terms, if there are multiple branches between them. For example, using the notations and demographic model in Fig. 1, we can estimate the total selection difference between East and West Africans as

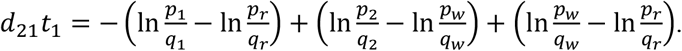

Let 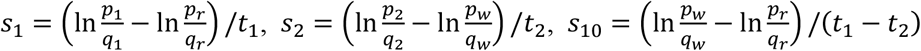, then we have

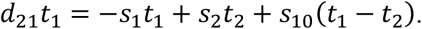

Therefore, we can represent the selection difference between paired populations as a combination of selection coefficients during different time periods. Using matrix algebra, we can represent the selection differences of all the paired populations in Fig. 1 as

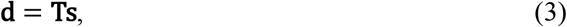

where

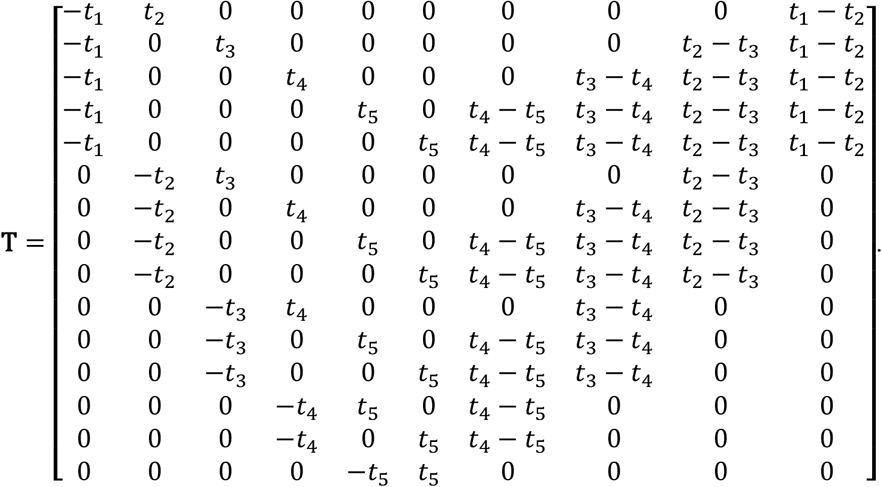

To obtain the optimal solution, we propose a probabilistic approach. Under neutral evolution (NE), we consider each estimated *s* as an independent random variable following a normal distribution with a mean of zero and a variance of *σ*^2^. For each solution with ten variables, the summation below follows a chi-square distribution with ten degrees of freedom:

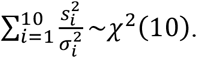

Therefore, we have Pr(|**s**|^2^ > |**s**_*a*_|^2^ | NE) ≥ Pr(|**s**|^2^ > |**s**_*b*_|^2^ | NE), if |**s**_*a*_|^2^ ≤ |**s**_*b*_|^2^ for solutions *a* and *b*. Here, |**s**|^2^ is the norm of a vector **s**. In other words, we can choose the most conservative solution with the least deviation from neutral evolution using a probabilistic approach. Thus, the optimal solution **s**\* is

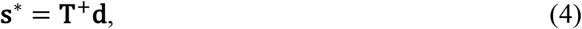

where **T**^+^ is the pseudo-inverse of **T**.

### Selective pressures over epochs

We applied our new approach with genotype data of worldwide populations from 17 publicly available datasets (Table S1). After data cleaning (Materials and Methods), we obtained 2346 individuals and grouped them into six population groups based on their geographic locations (Table S2). We also selected 52 SNPs in 19 genes for analysis due to their association with human pigmentation in published genome-wide association studies (GWAS) or phenotype prediction models (Table S3). We then used Eq. 2 with 31 SNPs not in strong linkage disequilibrium (*r*^2^ < 0.8) to estimate the total selection differences on human pigmentation (Materials and Methods). The maximum differences were observed between Europeans and the two African populations, while the minimum difference was observed between West and East Africans (Table 1). The estimated 95% confidence intervals (CI) indicate the difference between East and West Africans, as well as between Oceanians and East Asians, is likely due to genetic drift (Table 1). We further assessed the significance levels of the observed selection differences by randomly sampling 10,000 sets of 31 SNPs in the genome and obtained the empirical distributions of population differences (Fig. 1). The differences from random sets of SNPs were close to zero, whereas those from SNPs associated with human pigmentation were significantly departure from zero (Fig. 1). This suggests that most population differences on SNPs associated with human pigmentation are possibly contributed by natural selection and not confounded by genetic drift.

**Table 1.** Selection differences with 95% CI associated with human pigmentation between populations (× 10^−3^/generation).

|  | WAF | EAF | OCN | EUR | NAS | EAS |
| --- | --- | --- | --- | --- | --- | --- |
| **WAF** | 0 | -3.527<br>$\pm 3.745$ | -17.402<br>$\pm 6.623$ | -33.992<br>$\pm 5.425$ | -24.397<br>$\pm 6.359$ | -20.052<br>$\pm 6.084$ |
| **EAF** | 3.527<br>$\pm 3.745$ | 0 | -17.614<br>$\pm 8.391$ | -38.668<br>$\pm 6.900$ | -26.484<br>$\pm 8.085$ | -20.976<br>$\pm 7.753$ |
| **OCN** | 17.402<br>$\pm 6.623$ | 17.614<br>$\pm 8.391$ | 0 | -27.368<br>$\pm 8.695$ | -11.534<br>$\pm 9.078$ | -4.373<br>$\pm 8.511$ |
| **EUR** | 33.992<br>$\pm 5.425$ | 38.668<br>$\pm 6.900$ | 27.368<br>$\pm 8.695$ | 0 | 19.793<br>$\pm 9.400$ | 28.747<br>$\pm 8.907$ |
| **NAS** | 24.397<br>$\pm 6.359$ | 26.484<br>$\pm 8.085$ | 11.534<br>$\pm 9.078$ | -19.793<br>$\pm 9.400$ | 0 | 11.018<br>$\pm 8.741$ |
| **EAS** | 20.052<br>$\pm 6.084$ | 20.976<br>$\pm 7.753$ | 4.373<br>$\pm 8.511$ | -28.747<br>$\pm 8.907$ | -11.018<br>$\pm 8.741$ | 0 |

Populations are abbreviated as follows: WAF, West Africans; EAF, East Africans; OCN, Oceanians; EUR, Europeans; NAS, North Asians; EAS, East Asians.

We then solved the linear system (Eq. 4) with the observed selection differences on human pigmentation (Table 1). Our estimate shows that the modern European lineage had the largest selective pressure 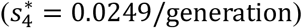 on light pigmentation than the other branches (Fig. 1), suggesting that recent natural selection favored light pigmentation in Europeans. Recent studies using ancient DNA could support our observation of recent directional selection in Europeans [18, 41]. Our results also reveal the selective pressure on light pigmentation in the ancestral population of Europeans and East Asians 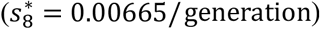. This shared selection is also supported by other studies, revealing that *ASIP, BNC2*, and *KITLG* were under directional selection before the divergence of ancestral Europeans and East Asians [16, 34]. We further applied SLiM 2 [42] to examine whether the optimal solution could reproduce the observed selection differences (Table 1). We set up a human demographic model according to previous studies and used the optimal solution as selection coefficients during different periods (Materials and Methods). The simulated selection differences were close to the data and little affected by the initial frequency of the beneficial allele (Fig. 3). This also illustrates that though we assume genic selection, our model could approximate genotypic selection in diploids (Materials and Methods).

**Fig. 2.**
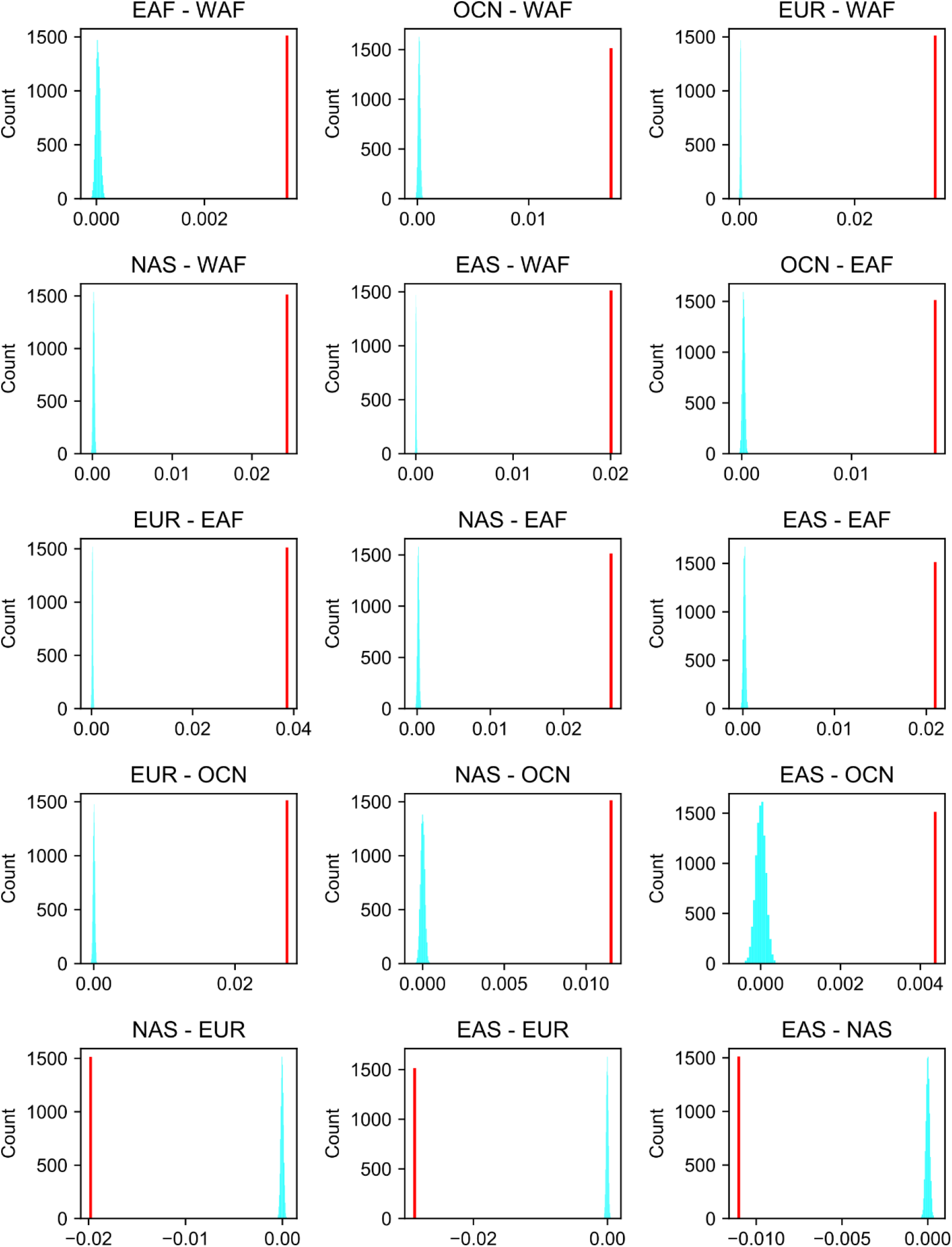
Significance levels of the selection differences associated with human pigmentation between populations. The blue bars are the empirical distributions of population differences using 10,000 random sets of 31 SNPs in the genomes. The red lines are the selection differences between populations using 31 SNPs associated human pigmentation (Materials and Methods). Population abbreviations: WAF, West Africans; EAF, East Africans; OCN, Oceanians; EUR, Europeans; NAS, North Asians; EAS, East Asians.

**Fig. 3.**
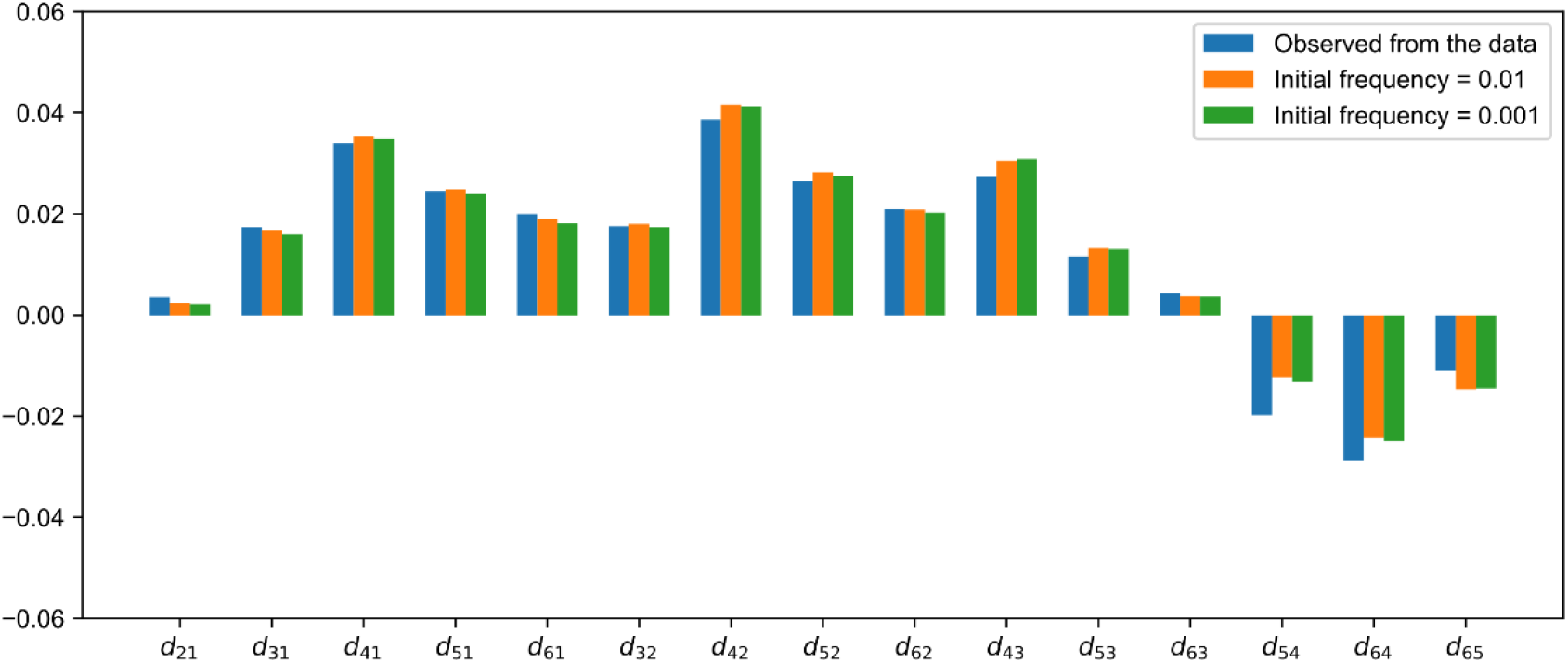
Comparisons between selection differences from simulation and the data. The selection differences are: *d*_21_, differences between East Africans and West Africans; *d*_31_, differences between Oceanians and West Africans; *d*_41_, differences between Europeans and West Africans; *d*_51_, differences between North Asians and West Africans; *d*_61_, differences between East Asians and West Africans; *d*_32_, differences between Oceanians and East Africans; *d*_42_, differences between Europeans and East Africans; *d*_52_, differences between North Asians and East Africans; *d*_62_, differences between East Asians and East Africans; *d*_43_, differences between Europeans and Oceanians; *d*_53_, differences between North Asians and Oceanians; *d*_63_, differences between East Asians and Oceanians; *d*_54_, differences between North Asians and Europeans; *d*_64_, differences between East Asians and Europeans; *d*_65_, differences between East Asians and North Asians.

### Selection differences between populations

We also separately quantified selection differences of individual SNPs associated with human pigmentation (Table S4) using Eq. 1. Ten SNPs were removed because of their low derived allele frequencies among populations in our data (Materials and Methods). Statistical tests suggest that selective pressures in many loci differed significantly between populations (*p* < 0.05). The remaining 42 SNPs were categorized into five groups (Fig. 4).

**Fig. 4.**
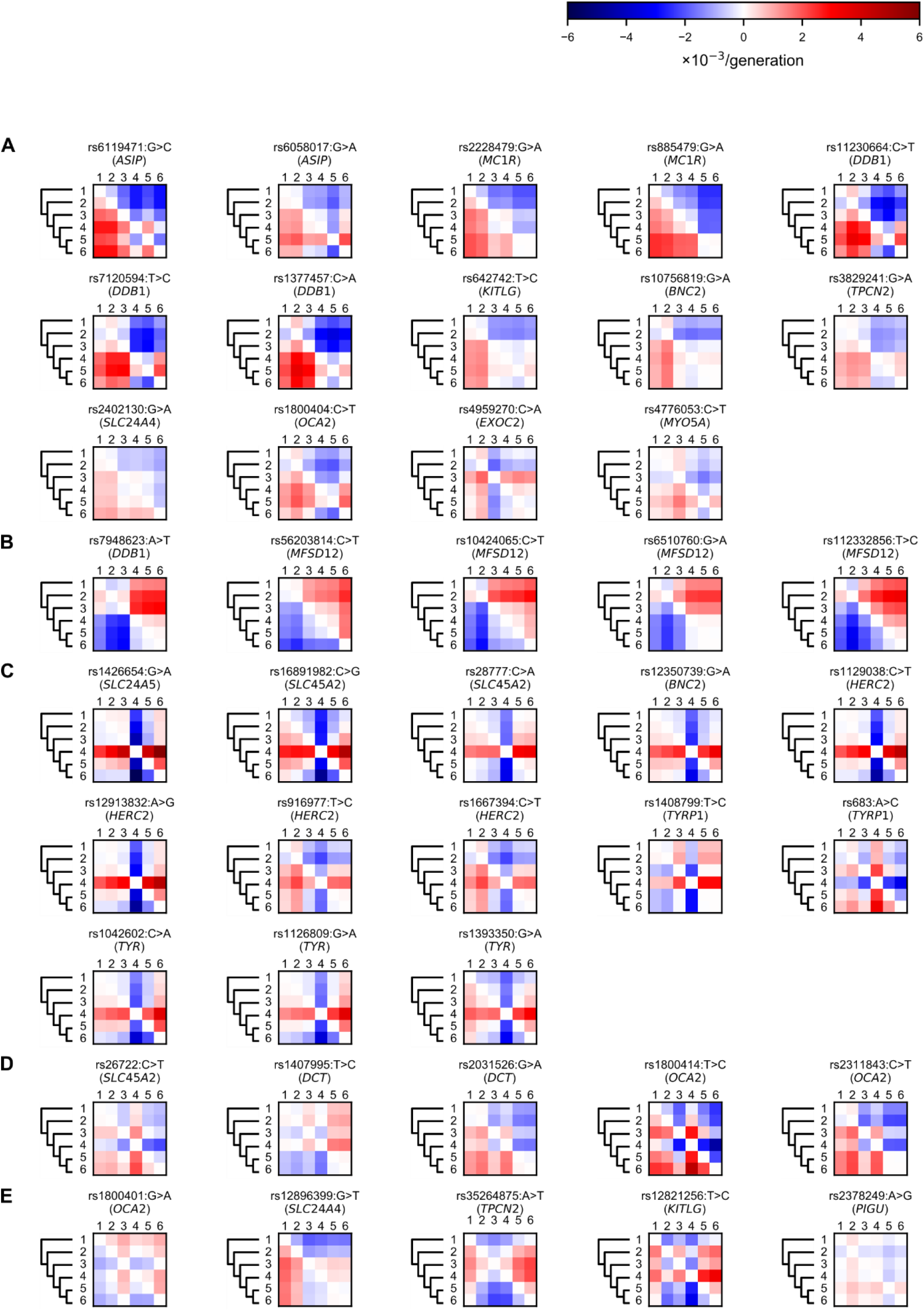
Selection differences in individual loci between populations. We used Eq. 1 to quantify the selection differences of 42 SNPs associated with human pigmentation, and categorized them into five kinds of patterns: (A) Derived alleles affected by Eurasian-shared selection; (B) Derived alleles affected by African-specific selection; (C) Alleles affected by European-specific selection; (D) Alleles affected by Asian-specific selection; and (E) Others. Red color (positive numbers) indicates selective pressures of populations in rows are larger than those in columns; blue color (negative numbers) indicates selective pressures of populations in rows are smaller than those in columns. Populations are abbreviated as follows: 1, West Africans; 2, East Africans; 3, Oceanians; 4, Europeans; 5, North Asians; 6, East Asians

In the first group, derived alleles may be affected by Eurasian-shared selection (Fig. 4A). Among these SNPs, rs6119471 (*ASIP*) has large selection difference between Eurasians and Africans (Table S4). The derived allele of rs6119471 (*ASIP*) is almost fixed across Eurasians but maintains a low frequency in Africans (Fig. S2). Recent studies applied this SNP to predict dark/non-dark pigmentation phenotype in humans [43]. This may be explained by different selective pressures on this SNP among populations. Other SNPs, such as rs642742 (*KITLG*) and rs10756819 (*BNC2*), exhibit non-significant differences, suggesting that selection may contribute less to the diversity of these SNPs.

Our results also indicate that two SNPs in *MC1R* (rs2228479 and rs885479) largely differ between Eurasians and Africans (Table S4). Previous studies used variants in *MC1R* to solve a long-standing puzzle, regarding whether light pigmentation in low UVR areas is caused by directional selection or the relaxation of selective pressures [18, 44, 45]. The relaxation of selective pressures would suggest that the diversity of *MC1R* variants increased in Eurasians due to the lack of selective constraints. In this scenario, the genetic diversity of *MC1R* variants could be largely attributed to genetic drift. In contrast, directional selection would suggest that *MC1R* variants were under positive selection in Eurasians. In this scenario, genetic drift cannot explain the genetic divergence of *MC1R* between Africans and Eurasians. Our statistical results show that the divergences of rs2228479 and rs885479 between Eurasians and Africans are highly significant departure from neutral evolution (Table S4), suggesting that directional selection is the more likely explanation. Experimental evidence suggests that the derived allele of rs2228479 could cause lower affinity for alpha-melanocyte stimulating hormone than the ancestral allele [46]. Another study showed that the derived allele of rs885479 carries a lower risk of developing freckles and severe solar lentigines than the ancestral allele in East Asians [47]. These studies revealed the potential roles of these *MC1R* variants in pigmentation phenotypes.

In the second group, derived alleles may be affected by African-specific selection (Fig. 4B). All these derived alleles are in/near two genes (*DDB1* and *MFSD12*) and were recently associated with human dark pigmentation [48]. The previous study [48] did not find signals of positive selection at *MFSD12* using Tajima’s D or iHS. Our method [36] show that these SNPs in *MFSD12* differ significantly between Africans and Eurasians—possible signals of directional selection (Table S4). From the first and second groups, we can observe that directional selection not only affects derived alleles associated with light pigmentation in Eurasians, but also influences derived alleles associated with dark pigmentation in Africans. This observation suggests that human pigmentation is under directional selection with diversifying orientations among different populations. Thus, the previous view that the dark pigmentation in Africans is the result of purifying selection on ancestral alleles is incomplete [45]. Moreover, these new loci illustrate that we still have little understanding of the genetic diversity within African populations, and more research in this area is needed [49].

The third and fourth groups display European- and Asian-specific selection, respectively (Fig. 4C and 4D). One notable SNP is rs1426654 (*SLC24A5*), which had the largest selection difference between Europeans and East Asians in our study (0.005774/generation). Previous studies reported that this SNP is under strong directional selection in Europeans [10–15]. Another notable SNP is rs1800414 (*OCA2*), which had a large selection difference between East Asians and other populations. This reveals a potential role of rs1800414 (*OCA2*) on light pigmentation in East Asians. Several studies have suggested directional selection on this SNP in East Asians [50, 51]. These large selection differences indicate the significant contributions of these SNPs to light pigmentation in Europeans and East Asians, respectively. Other SNPs in these groups also support the hypothesis that recent natural selection for light pigmentation independently occurred in Europeans and Asians since they diverged [14, 50, 51]. Interestingly, Oceanians comprise both African-specific (*DDB1*) and Asian-specific (*OCA2*) selection. However, due to limited sample size of Oceanians in our data from publicly available resources (Table S2), it should be cautious to interpret these results. Thus, it would be helpful to analyze larger datasets of Oceanians to confirm our observation.

The last group includes the five remaining SNPs (Fig. 4E), which exhibit specific selection differences between limited population pairs. Among them, the derived allele of rs1800401 (OCA2) and the ancestral allele of rs12896399 (*SLC24A4*) are both associated with dark pigmentation (Table S2). Only rs12896399 (*SLC24A4*) differs significantly between West Africans and Eurasians (Table S4). This may be a selection signal associated with dark pigmentation in West Africans, again indicating possible genetic diversity within African populations. We note that rs35264875 (*TPCN2*) and rs12821256 (*KITLG*) might be affected by selection in both East Africans and Europeans. A recent study showed that rs12821256 might have large effect on the skin pigmentation in South Africans [52]. The other two SNPs, rs3829241 (*TPCN2*) and rs642742 (*KITLG*), also differ between Eurasians and Africans (Fig. 4A). These similar patterns of *TPCN2* and *KITLG* might suggest some connection between them.

Compared with previous studies [10–18], our study has three advantages. First, our approach considers the fluctuation of selective pressures that was ignored by previous studies. Our results provide more information about the dynamics of selective pressures during human evolution. Second, we summarize selective pressures based on multiple human pigmentation genes (Eq. 2), while previous studies usually tested selection signals in individual human pigmentation genes. Moreover, we simultaneously interpret selective pressures in multiple populations, whereas previous studies separately investigated selection signals in single population. Third, we do not need to assume population continuity as in those ancient DNA studies [18, 41], because our study is based on genetic data from only present-day populations.

We note that our investigation has several limitations. First, our model is based on the infinite population size model. The limited sample size would affect our results, therefore, we grouped populations into a large population group based on their geographic locations to mitigate the effect of sample size. Analysis of data with larger sample size could improve our estimate, as more and more genomic datasets become available. Second, although we chose the solution that deviates least from neutral evolution as the optimal solution, we cannot exclude the possibility of other solutions. This reflects the difficulty of analyzing historical selective pressures, which is a well-recognized challenge in population genetics [53]. Our solution provides a first step toward resolving the dynamics of selection in the evolution of human pigmentation. This solution may be improved by combining both ancient and modern human genetic data, as well as by using a Bayesian framework with varied selection coefficients among loci for inference. Adding more population groups would also possibly improve the solution, because this would provide more constraints in the linear system (Eq. 4). Third, our results may be affected by a severe bottleneck. A recent study [53] suggests a more severe Out-of-Africa bottleneck in human evolutionary history than in the model used in our simulation. This would probably reduce the selection differences between Eurasians and Africans, leading to an underestimation of selective pressures. Fourth, our results may also be affected by population migration and sub-structure. We used knowledge from previous studies, principle component analysis and *F*_3_ test to rigorously prune potential admixed populations, including South Asians, Central Asians, the Middle East People and Americans. Removing these populations would lose information of selective pressures on human pigmentation in these lineages; however, as a first step to explore the historical selective pressures in the evolution of human pigmentation, we focused more on reducing the bias induced by population admixture. New methods explicitly accounting for population admixture would be helpful to provide more comprehensive view on the dynamics of selective pressures during the evolution of human pigmentation. Besides, we demonstrate that our estimate provides lower bounds of selection differences on human pigmentation when migration or sub-structure exists (Text S1). Finally, our candidate SNPs may be biased. For example, our results indicate small genetic differences on human pigmentation between these two populations (Table 1), while recent studies [52] suggest Oceanians are darker than East Asians in skin pigmentation using melanin index. One possible reason is that some Oceanian-specific or East-Asian-specific SNPs are missing. This is because we selected candidates based on results from published GWAS or phenotype prediction models, and most of these studies used samples with European ancestry [55]. More studies on non-European populations could resolve this missing diversity and enhance our knowledge on the evolution of human pigmentation.

To summarize, we extended an established method [36] to dissect dynamics of selective pressures over epochs. Our study provides the first attempt to resolve time-varied selective pressures in the evolution of human pigmentation. The fluctuation of selective pressures is an important factor in evolution [53], but few studies have revealed time-varied selective pressures in human populations. Our study also provides information on differences of selective pressures between distinct population groups. Further studies are in progress to verify our present views on the evolution of human pigmentation.

## Materials and Methods

### Data preparation

Seventeen datasets [40, 56–71] containing genotype data from worldwide human populations were obtained from the listed resources (Table S1). After downloading, all the genotype data were liftovered to genomic coordinates using the Human Reference Genome hg19. A merged dataset containing 6531 individuals was obtained after removing duplicated and related individuals. After merging, SNPs with call rate less than 0.99 or individuals with call rate less than 0.95 were removed. SNPs in strong linkage disequilibrium were further removed by applying a window of 200 SNPs advanced by 25 SNPs and an *r*^2^ threshold of 0.8 (--indep-pairwise 200 25 0.8) in PLINK 1.7 [72]. The remaining 61,597 SNPs were used for further analysis. In order to mitigate the bias induced by population migration, potential admixed populations, such as the Middle East People and South Asians, were excluded according to previous studies [40, 56–71], principal component analysis (PCA) using SMARTPCA (version: 13050) from EIGENSOFT (version: 6.0.1) [73, 74], and *F*_3_ test using ADMIXTOOLS (version: 3.0) [75]. Finally, 2346 individuals were obtained and divided into six groups according to their geographic regions for further analysis. These groups are West Africans, East Africans, Oceanians, Europeans, North Asians and East Asians. The PCA plot (Fig. S1) shows that these 2346 individuals were properly separated into six population groups.

**Fig. S1.**
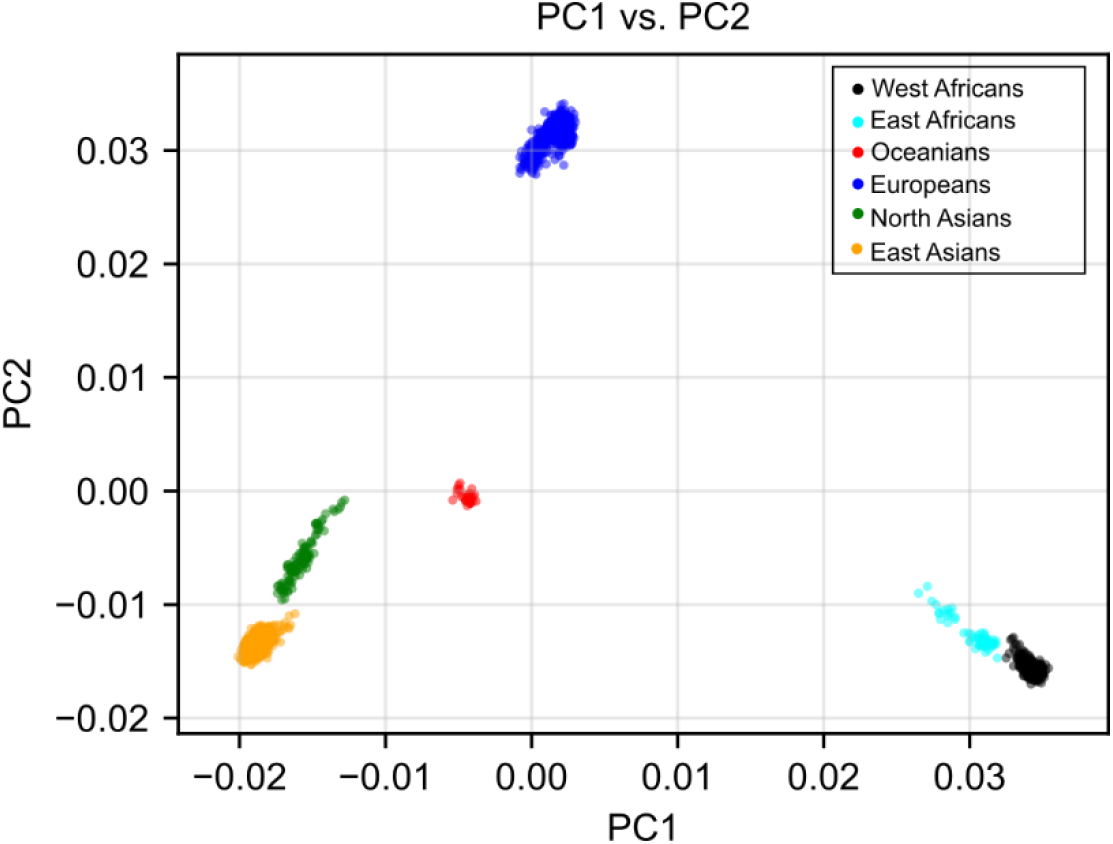
PCA plots of 2346 samples.

### Data imputation

Genotypes of 19 human pigmentation genes with 500-kb flanking sequences on both sides were obtained from the genotype datasets. Haplotype inference and genotype imputation were performed on the selected genotypes using BEAGLE 4.1 [76, 77] with 1000 Genomes phase 3 haplotypes as the reference panel. During phasing and imputation, the effective population size was assumed to be 10,000 (*N_e_* = 10000), and the other parameters were set to the default values. Ten SNPs (rs1110400, rs11547464, rs12203592, rs1800407, rs1805005, rs1805006, rs1805007, rs1805008, rs1805009, rs74653330) were removed because of their low derived allele frequencies in our datasets after imputation (Fig. S2). Because rs12203592 (*IRF4*) was removed, 18 genes with the remaining 42 SNPs were used for further analysis.

**Fig. S2.**
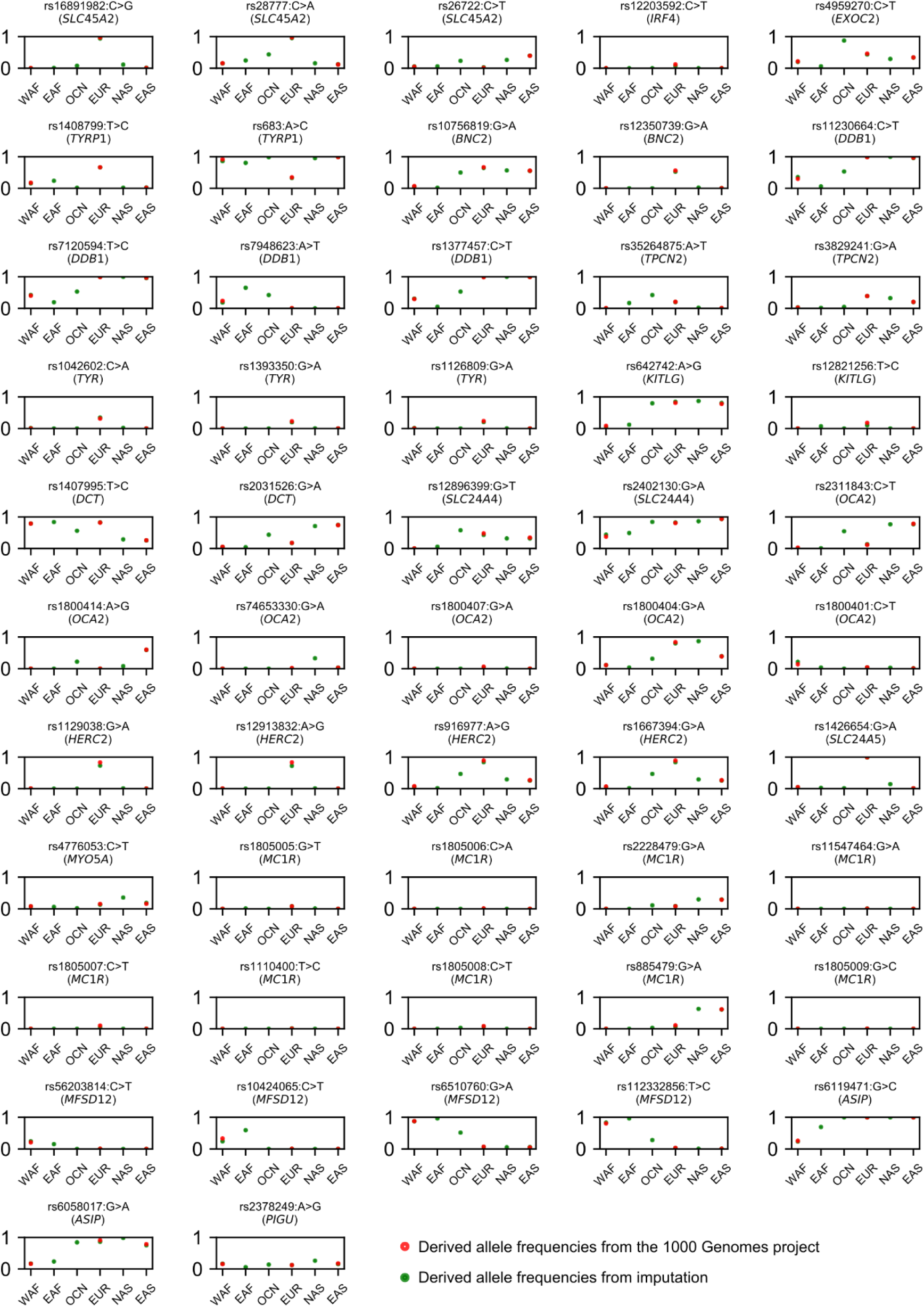
Derived allele frequencies of candidate SNPs from imputation and the 1000 Genomes project. Population abbreviations: WAF, West Africans; EAF, East Africans; OCN, Oceanians; EUR, Europeans; NAS, North Asians; EAS, East Asians.

### Estimating selection differences between populations and selective pressures over epochs

We used Eq. 1 to estimate the selection differences of the remaining 42 SNPs. We then used Eq. 2 and selected 31 SNPs not in strong linkage disequilibrium (*r*^2^ < 0.8) as well as known phenotypes to estimate the total selection differences on human pigmentation between populations. These SNPs were: rs3829241, rs56203814, rs916977, rs1800414, rs10424065, rs6119471, rs1408799, rs11230664, rs1407995, rs4959270, rs1800401, rs2378249, rs1042602, rs12350739, rs6058017, rs12821256, rs1393350, rs1426654, rs642742, rs6510760, rs1129038, rs2228479, rs35264875, rs12896399, rs26722, rs16891982, rs885479, rs28777, rs1800404, rs10756819, rs2402130. To dissect selective pressures over epochs, we applied Eq. 4 with the total selection differences from the selected 31 SNPs and the divergence times shown in Fig. 1.

### Reproducing the observed selection differences from the optimal solution

We used SLiM 2 (version: 2.6) [42] to simulate a demographic model of human evolution (Fig. S3) to examine whether the optimal solution could reproduce the observed selection differences. We varied the initial frequency of the beneficial allele with 0.001 and 0.01. We divided the optimal solution by 31 to obtain the average selection coefficient for each SNP, because we used 31 SNPs to estimate the total selection differences on human pigmentation (Text S2). We used the effective population size of each population estimated by previous studies [78, 79]. We set both the mutation rate and the recombination rate to 1 × 10^−8^/generation/site. In each run, we simulated a fragment with 10^6^ base pairs, and set the 50,000th site under selection. We repeated each set of parameters more than 10,000 times, and analyzed those results in which beneficial alleles were not fixed or lost in all the populations. We compared the average selection differences from simulation with the observed selection differences.

All the simulations were performed in Digital Ocean (https://cloud.digitalocean.com/) Optimized Droplets. The information of these droplets is as follows: CPU, Intel^®^ Xeon^®^ Platinum 8168 Processor; Random-access memory, 64 GB; Operating system, Ubuntu 16.04.4 x64.

**Fig. S3.**
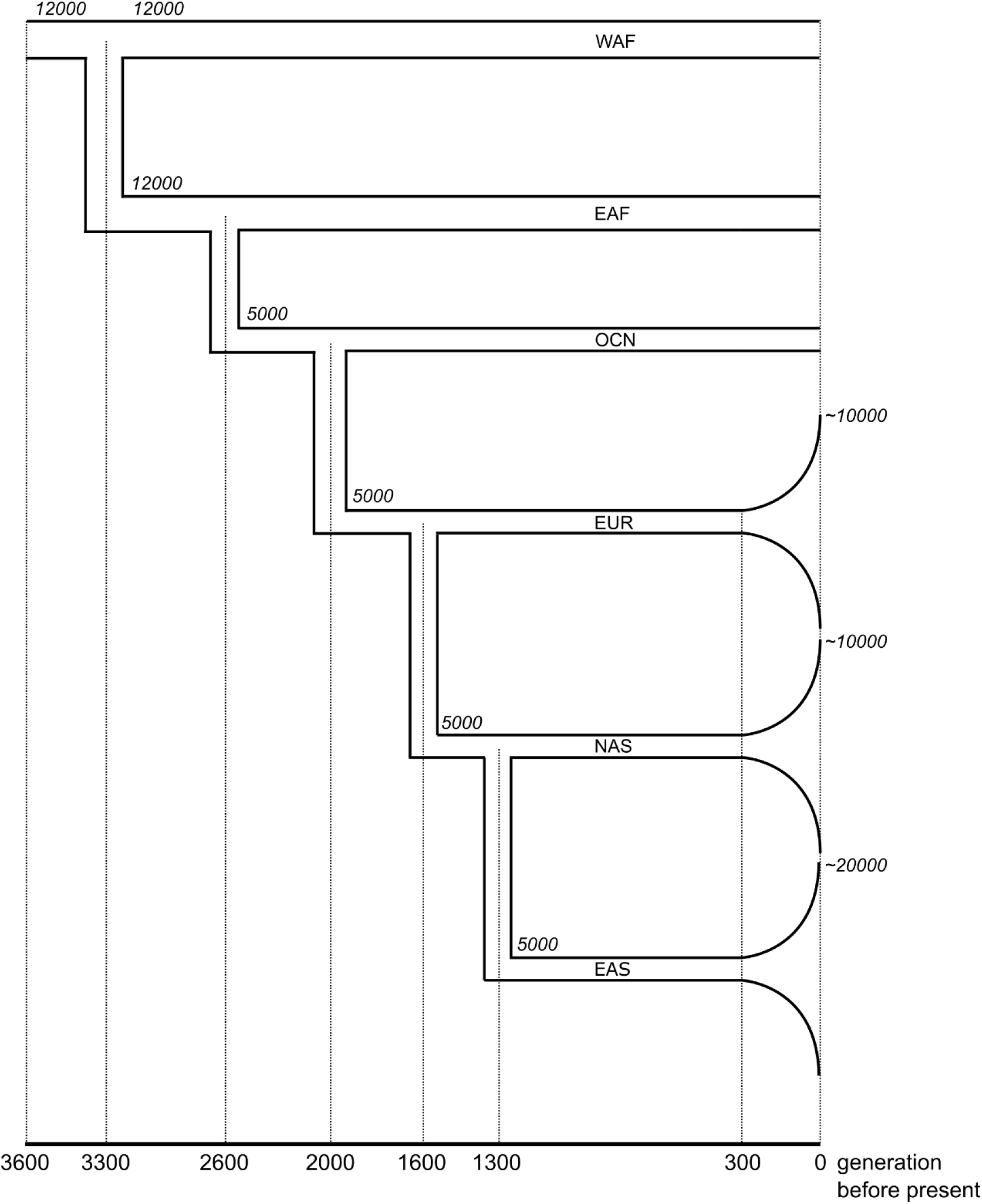
The demography model for simulation. The italic numbers indicate population sizes at different time periods. Population abbreviations: WAF, West Africans; EAF, East Africans; OCN, Oceanians; EUR, Europeans; NAS, North Asians; EAS, East Asians.

We noticed that the selection coefficient in SLiM 2 measures differences in fitness between genotypes instead of alleles. We can transform the selection coefficient of genotypes into that of alleles as follows. Let the fitness of the ancestral allele *A* be 1, and the relative fitness of the derived allele *a* is *e^s^*. When *s* is close to 0, we can approximate *e^s^* as 1 + *s* using the Taylor series. The fitness of genotype *aa* becomes (1 + *s*)^2^ = 1 + 2*s* + *s*^2^ ≈1 + 2*s*, and the fitness of genotype *Aa* is 1 + *s* = 1 + 0.5*s′*. If *s′* is the selection coefficient in SLiM 2, then *s′* = 2*s*; and the dominance coefficient becomes 0.5.

## Supporting Information

**S1 Table. Data resources.**

**S2 Table. Population information.**

**S3 Table. Candidate SNPs.**

**S4 Table. Selection differences on the selected 42 SNPs.**

**S1 Text. The effects of migration and substructure.**

**S2 Text. A sample script for simulation with SLiM 2.**

## Acknowledgements

This work was supported by grants from National Natural Science Foundation of China (91331109 to Y.H.; 31271338 and 31330038 to L.J.; 31322030 and 91331108 to S.W.). Y.H. is also grateful for support from the SA-SIBS scholarship program and the Youth Innovation Promotion Association of Chinese Academy of Science (2012216). L.J. was also supported by the Shanghai Leading Academic Discipline Project (B111) and the Center for Evolutionary Biology at Fudan University. S.W. was also awarded by the National Thousand Young Talents Award, the Max Planck-CAS Paul Gerson Unna Independent Research Group Leadership Award, and open projects from the State Key Laboratory of Genetic Engineering at Fudan University. X.H. is grateful to Dr. Minxian Wang, Dr. Yuchen Wang, Dr. Haiyi Lou and Dr. Lin Tang for comments on the manuscript.

## Author Contribution

X.H. and Y.H. designed the study. X.H. collected and simulated the data. X.H. and Y.H. developed the model, analyzed the data and wrote the manuscript. S.W. and L.J. revised the manuscript.

## Supporting information

**Table S1.** Data resources

| Dataset | Sample | Resource | Reference |
| --- | --- | --- | --- |
| 1KG | 2504 | http://www.1000genomes.org/ | 66 |
| HapMap3 | 1397 | http://www.sanger.ac.uk/resources/downloads/human/hapmap3.html | 67 |
| Jew | 466 | http://evolbio.ut.ee/jew/ | 57 |
| Afghan | 24 | http://evolbio.ut.ee/afghan/ | 58 |
| Sakha | 40 | http://evolbio.ut.ee/sakha/ | 59 |
| Balkan | 70 | http://evolbio.ut.ee/balkan/ | 60 |
| HGDP | 1043 | ftp://ftp.cephb.fr/hgdp_supp1/ | 56 |
| India | 142 | http://evolbio.ut.ee/india/ | 61 |
| Ethiopian | 235 | http://mega.bioanth.cam.ac.uk/data/Ethiopia | 62 |
| Malta | 85 | http://evolbio.ut.ee/malta/ | 63 |
| Saqqaq | 197 | http://evolbio.ut.ee/saqqaq/ | 64 |
| SGVP | 268 | http://phg.nus.edu.sg/StatGen/public_html/SGVP/download.html | 65 |
| NorthernEurasian | 369 | Request from the authors | 68 |
| Caucasus | 204 | http://evolbio.ut.ee/caucasus/ | 69 |
| Turkic | 322 | http://evolbio.ut.ee/turkic/ | 70 |
| Andamanese | 10 | https://www.ebi.ac.uk/ena/data/view/PRJEB11455 | 40 |
| EGDP | 483 | http://evolbio.ut.ee/CGgenomes.html | 71 |

**Table S2.** Population information

| Data resource | Population group | Population abbreviation | Sample size |
| --- | --- | --- | --- |
| Ethiopian | East Africans | ANU | 21 |
| Ethiopian | East Africans | GUM | 15 |
| Ethiopian | East Africans | SUD | 23 |
| 1KG | East Asians | CHN | 208 |
| 1KG | East Asians | DAI | 93 |
| 1KG | East Asians | JAP | 104 |
| 1KG | East Asians | KHV | 96 |
| EGDP | East Asians | BUM | 1 |
| EGDP | East Asians | DSN | 8 |
| EGDP | East Asians | IGO | 8 |
| EGDP | East Asians | LBO | 4 |
| EGDP | East Asians | LUZ | 2 |
| EGDP | East Asians | MUR | 8 |
| EGDP | East Asians | VIZ | 2 |
| EGDP | East Asians | VTN | 10 |
| HapMap3 | East Asians | CHN | 152 |
| HapMap3 | East Asians | JAP | 16 |
| HGDP | East Asians | CHN | 44 |
| HGDP | East Asians | DAI | 10 |
| HGDP | East Asians | JAP | 28 |
| HGDP | East Asians | LAH | 8 |
| HGDP | East Asians | MIA | 10 |
| HGDP | East Asians | NAX | 9 |
| HGDP | East Asians | SHE | 10 |
| HGDP | East Asians | TUJ | 10 |
| HGDP | East Asians | YIZ | 10 |
| SGVP | East Asians | CHN | 96 |
| 1KG | Europeans | CEU | 99 |
| 1KG | Europeans | FIN | 99 |
| 1KG | Europeans | GBR | 91 |
| Jew | Europeans | BEL | 9 |
| Jew | Europeans | HNG | 20 |
| Jew | Europeans | LIT | 10 |
| Jew | Europeans | RMN | 14 |
| Jew | Europeans | RUS | 2 |
| EGDP | Europeans | ALB | 3 |
| EGDP | Europeans | BEL | 4 |
| EGDP | Europeans | COS | 4 |
| EGDP | Europeans | CRO | 4 |
| EGDP | Europeans | EST | 6 |
| EGDP | Europeans | FIN | 3 |
| EGDP | Europeans | GER | 3 |
| EGDP | Europeans | HNG | 2 |
| EGDP | Europeans | ING | 3 |
| EGDP | Europeans | KAR | 3 |
| EGDP | Europeans | KOM | 2 |
| EGDP | Europeans | LAT | 3 |
| EGDP | Europeans | LIT | 3 |
| EGDP | Europeans | MOL | 2 |
| EGDP | Europeans | MRD | 3 |
| EGDP | Europeans | POL | 4 |
| EGDP | Europeans | RUS | 7 |
| EGDP | Europeans | SWE | 2 |
| EGDP | Europeans | UKR | 7 |
| EGDP | Europeans | VEP | 4 |
| HapMap3 | Europeans | CEU | 24 |
| Balkan | Europeans | BOS | 14 |
| Balkan | Europeans | KSV | 9 |
| Balkan | Europeans | MAC | 14 |
| Balkan | Europeans | MNT | 14 |
| Balkan | Europeans | SER | 18 |
| HGDP | Europeans | FRE | 28 |
| HGDP | Europeans | ORC | 15 |
| HGDP | Europeans | RUS | 25 |
| Malta | Europeans | EST | 14 |
| Malta | Europeans | RUS | 1 |
| NorthernEurasian | Europeans | SLV | 25 |
| Turkic | Europeans | GAG | 12 |
| Turkic | Europeans | GER | 13 |
| Turkic | Europeans | KAR | 15 |
| Turkic | Europeans | RUS | 33 |
| Turkic | Europeans | VEP | 11 |
| EGDP | North Asians | EVK | 13 |
| EGDP | North Asians | EVN | 8 |
| EGDP | North Asians | NGA | 2 |
| EGDP | North Asians | SAK | 7 |
| EGDP | North Asians | YAK | 1 |
| Sakha | North Asians | DOL | 3 |
| Sakha | North Asians | EVN | 8 |
| Sakha | North Asians | YAK | 1 |
| HGDP | North Asians | YAK | 22 |
| Malta | North Asians | DOL | 1 |
| Malta | North Asians | EVE | 2 |
| Saqqaq | North Asians | DOL | 4 |
| Saqqaq | North Asians | EVE | 15 |
| Saqqaq | North Asians | NGA | 13 |
| Saqqaq | North Asians | YUK | 3 |
| Turkic | North Asians | EVE | 3 |
| Turkic | North Asians | EVN | 3 |
| Turkic | North Asians | NGA | 2 |
| Turkic | North Asians | YAK | 3 |
| EGDP | Oceanians | KOI | 3 |
| EGDP | Oceanians | KOS | 3 |
| HGDP | Oceanians | PAP | 16 |
| Andamanese | Oceanians | AND | 10 |
| 1KG | West Africans | ESN | 99 |
| 1KG | West Africans | GWD | 112 |
| 1KG | West Africans | MSL | 85 |
| 1KG | West Africans | YOR | 108 |
| HapMap3 | West Africans | YOR | 46 |
| HGDP | West Africans | MND | 22 |
| HGDP | West Africans | YOR | 21 |

**Table S3.**
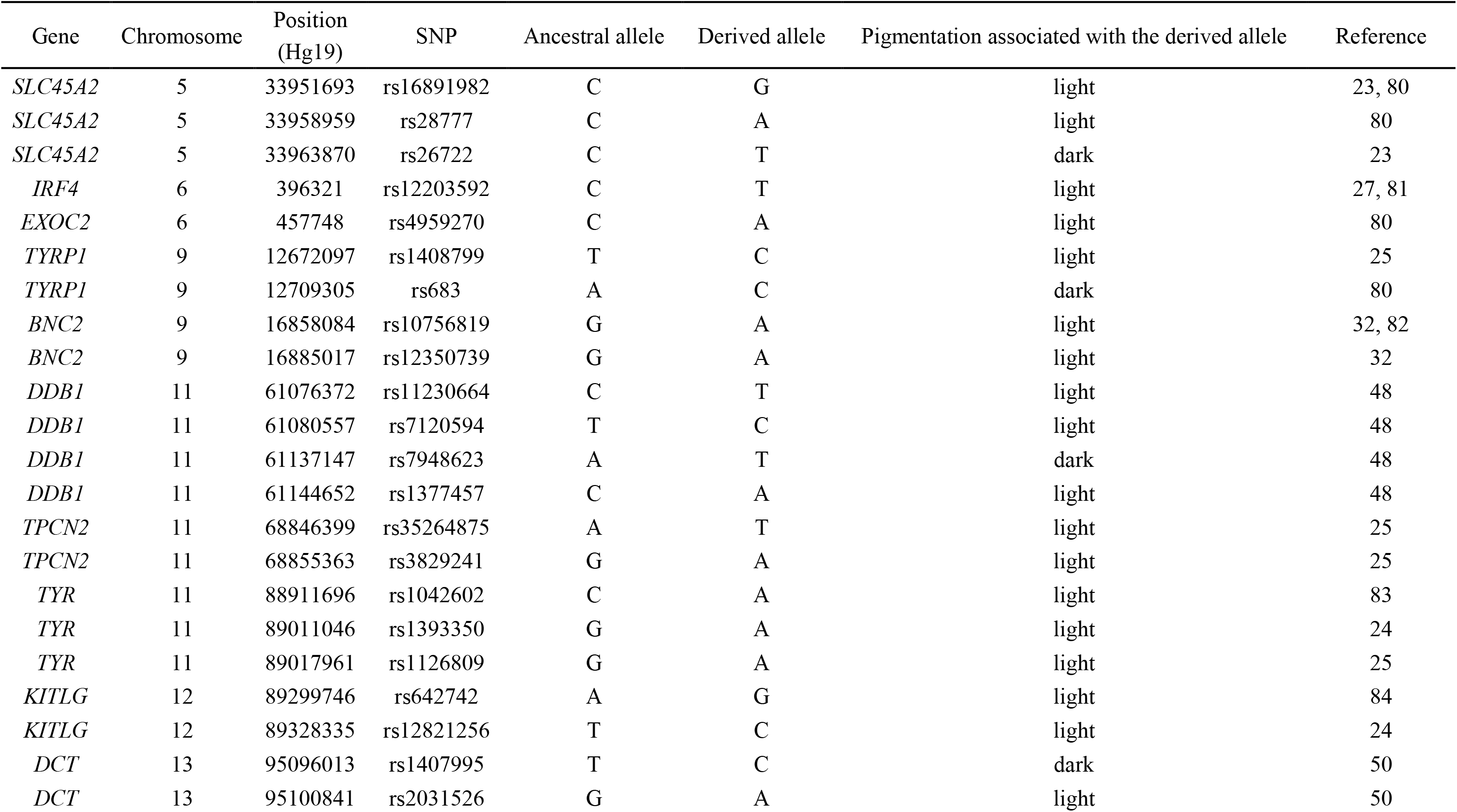
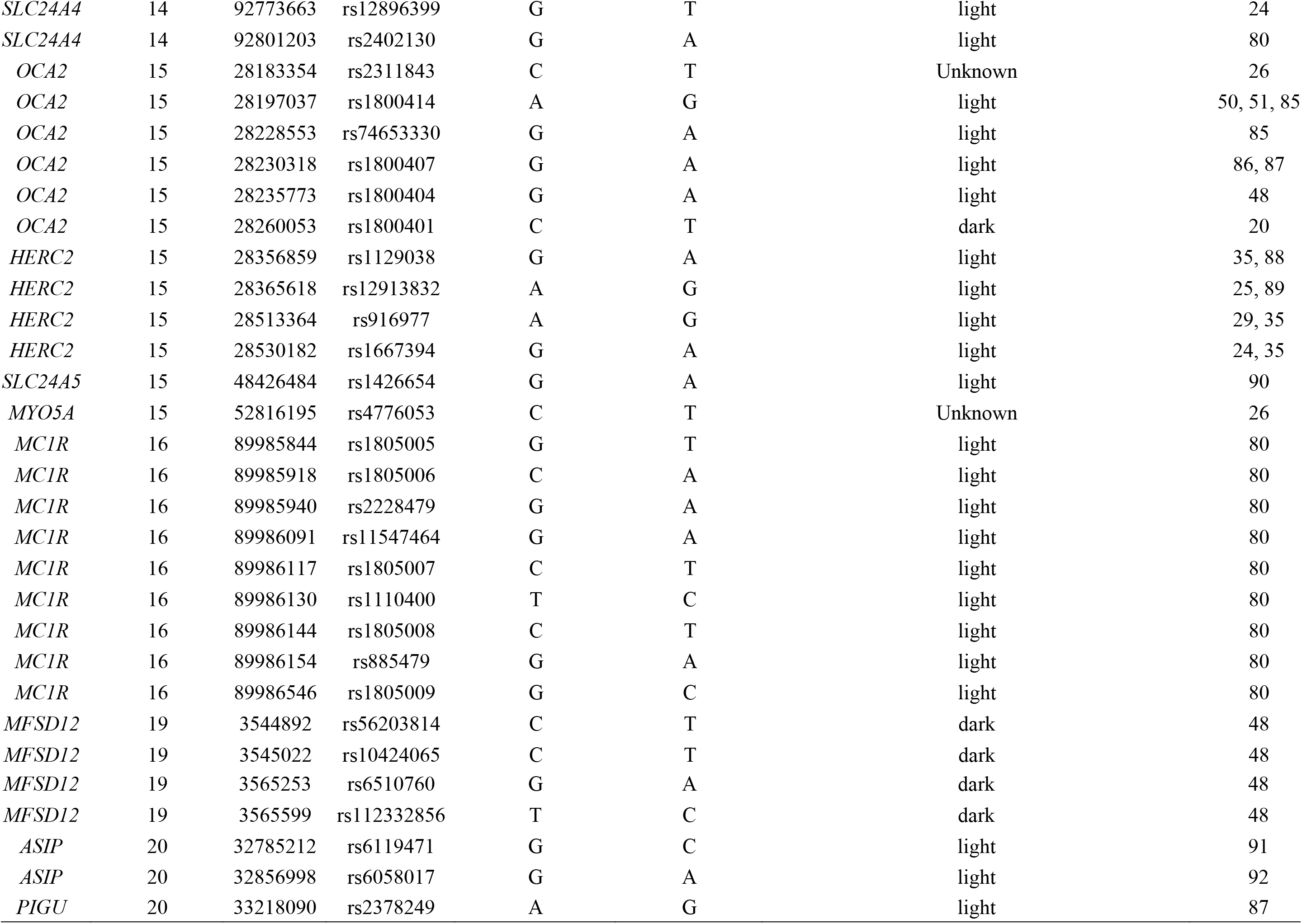
Candidate SNP information

| Gene | Chromosome | Position (Hg19) | SNP | Ancestral allele | Derived allele | Pigmentation associated with the derived allele | Reference |
| --- | --- | --- | --- | --- | --- | --- | --- |
| *SLC45A2* | 5 | 33951693 | rs16891982 | C | G | light | 23, 80 |
| *SLC45A2* | 5 | 33958959 | rs28777 | C | A | light | 80 |
| *SLC45A2* | 5 | 33963870 | rs26722 | C | T | dark | 23 |
| *IRF4* | 6 | 396321 | rs12203592 | C | T | light | 27, 81 |
| *EXOC2* | 6 | 457748 | rs4959270 | C | A | light | 80 |
| *TYRP1* | 9 | 12672097 | rs1408799 | T | C | light | 25 |
| *TYRP1* | 9 | 12709305 | rs683 | A | C | dark | 80 |
| *BNC2* | 9 | 16858084 | rs10756819 | G | A | light | 32, 82 |
| *BNC2* | 9 | 16885017 | rs12350739 | G | A | light | 32 |
| *DDB1* | 11 | 61076372 | rs11230664 | C | T | light | 48 |
| *DDB1* | 11 | 61080557 | rs7120594 | T | C | light | 48 |
| *DDB1* | 11 | 61137147 | rs7948623 | A | T | dark | 48 |
| *DDB1* | 11 | 61144652 | rs1377457 | C | A | light | 48 |
| *TPCN2* | 11 | 68846399 | rs35264875 | A | T | light | 25 |
| *TPCN2* | 11 | 68855363 | rs3829241 | G | A | light | 25 |
| *TYR* | 11 | 88911696 | rs1042602 | C | A | light | 83 |
| *TYR* | 11 | 89011046 | rs1393350 | G | A | light | 24 |
| *TYR* | 11 | 89017961 | rs1126809 | G | A | light | 25 |
| *KITLG* | 12 | 89299746 | rs642742 | A | G | light | 84 |
| *KITLG* | 12 | 89328335 | rs12821256 | T | C | light | 24 |
| *DCT* | 13 | 95096013 | rs1407995 | T | C | dark | 50 |
| *DCT* | 13 | 95100841 | rs2031526 | G | A | light | 50 |
| *SLC24A4* | 14 | 92773663 | rs12896399 | G | T | light | 24 |
| *SLC24A4* | 14 | 92801203 | rs2402130 | G | A | light | 80 |
| *OCA2* | 15 | 28183354 | rs2311843 | C | T | Unknown | 26 |
| *OCA2* | 15 | 28197037 | rs1800414 | A | G | light | 50, 51, 85 |
| *OCA2* | 15 | 28228553 | rs74653330 | G | A | light | 85 |
| *OCA2* | 15 | 28230318 | rs1800407 | G | A | light | 86, 87 |
| *OCA2* | 15 | 28235773 | rs1800404 | G | A | light | 48 |
| *OCA2* | 15 | 28260053 | rs1800401 | C | T | dark | 20 |
| *HERC2* | 15 | 28356859 | rs1129038 | G | A | light | 35, 88 |
| *HERC2* | 15 | 28365618 | rs12913832 | A | G | light | 25, 89 |
| *HERC2* | 15 | 28513364 | rs916977 | A | G | light | 29, 35 |
| *HERC2* | 15 | 28530182 | rs1667394 | G | A | light | 24, 35 |
| *SLC24A5* | 15 | 48426484 | rs1426654 | G | A | light | 90 |
| *MYO5A* | 15 | 52816195 | rs4776053 | C | T | Unknown | 26 |
| *MC1R* | 16 | 89985844 | rs1805005 | G | T | light | 80 |
| *MC1R* | 16 | 89985918 | rs1805006 | C | A | light | 80 |
| *MC1R* | 16 | 89985940 | rs2228479 | G | A | light | 80 |
| *MC1R* | 16 | 89986091 | rs11547464 | G | A | light | 80 |
| *MC1R* | 16 | 89986117 | rs1805007 | C | T | light | 80 |
| *MC1R* | 16 | 89986130 | rs1110400 | T | C | light | 80 |
| *MC1R* | 16 | 89986144 | rs1805008 | C | T | light | 80 |
| *MC1R* | 16 | 89986154 | rs885479 | G | A | light | 80 |
| *MC1R* | 16 | 89986546 | rs1805009 | G | C | light | 80 |
| *MFSD12* | 19 | 3544892 | rs56203814 | C | T | dark | 48 |
| *MFSD12* | 19 | 3545022 | rs10424065 | C | T | dark | 48 |
| *MFSD12* | 19 | 3565253 | rs6510760 | G | A | dark | 48 |
| *MFSD12* | 19 | 3565599 | rs112332856 | T | C | dark | 48 |
| *ASIP* | 20 | 32785212 | rs6119471 | G | C | light | 91 |
| *ASIP* | 20 | 32856998 | rs6058017 | G | A | light | 92 |
| *PIGU* | 20 | 33218090 | rs2378249 | A | G | light | 87 |

**Table S4.** Selection differences on the 42 selected SNPs

| Gene | SNP ID | Ancestral allele | Derived allele | Population1 | Population2 | Selection difference<br>(Population1 - Population2) | Std | p-value |
| --- | --- | --- | --- | --- | --- | --- | --- | --- |
| *SLC45A2* | rs16891982 | C | G | Europeans | NorthAsians | 0.003181 | 7.47E-04 | **2.10E-05** |
| *SLC45A2* | rs28777 | C | A | Europeans | NorthAsians | 0.003107 | 7.46E-04 | **3.10E-05** |
| *SLC45A2* | rs26722 | C | T | Europeans | NorthAsians | -0.001767 | 7.47E-04 | **0.017939** |
| *EXOC2* | rs4959270 | C | A | Europeans | NorthAsians | 4.12E-04 | 7.38E-04 | 0.576675 |
| *TYRP1* | rs1408799 | T | C | Europeans | NorthAsians | 0.00287 | 7.90E-04 | **2.82E-04** |
| *TYRP1* | rs683 | A | C | Europeans | NorthAsians | -0.002354 | 7.60E-04 | **0.001943** |
| *BNC2* | rs10756819 | G | A | Europeans | NorthAsians | 2.33E-04 | 7.37E-04 | 0.752243 |
| *BNC2* | rs12350739 | G | A | Europeans | NorthAsians | 0.002475 | 7.85E-04 | **0.00161** |
| *DDB1* | rs11230664 | C | T | Europeans | NorthAsians | -8.56E-04 | 0.001162 | 0.461178 |
| *DDB1* | rs7120594 | T | C | Europeans | NorthAsians | -5.17E-04 | 0.001172 | 0.658962 |
| *DDB1* | rs7948623 | A | T | Europeans | NorthAsians | 4.21E-04 | 0.001176 | 0.720668 |
| *DDB1* | rs1377457 | C | A | Europeans | NorthAsians | -7.42E-04 | 0.001165 | 0.524412 |
| *TPCN2* | rs35264875 | A | T | Europeans | NorthAsians | 0.00152 | 7.85E-04 | 0.052919 |
| *TPCN2* | rs3829241 | G | A | Europeans | NorthAsians | 1.79E-04 | 7.38E-04 | 0.808351 |
| *TYR* | rs1042602 | C | A | Europeans | NorthAsians | 0.002002 | 7.90E-04 | **0.011305** |
| *TYR* | rs1393350 | G | A | Europeans | NorthAsians | 0.0023 | 8.94E-04 | **0.010076** |
| *TYR* | rs1126809 | G | A | Europeans | NorthAsians | 0.002329 | 8.94E-04 | **0.009156** |
| *KITLG* | rs642742 | A | G | Europeans | NorthAsians | -1.94E-04 | 7.43E-04 | 0.794378 |
| *KITLG* | rs12821256 | T | C | Europeans | NorthAsians | 0.002653 | 0.001149 | **0.020951** |
| *DCT* | rs1407995 | T | C | Europeans | NorthAsians | 0.001528 | 7.38E-04 | **0.038476** |
| *DCT* | rs2031526 | G | A | Europeans | NorthAsians | -0.001525 | 7.38E-04 | **0.038867** |
| *SLC24A4* | rs12896399 | G | T | Europeans | NorthAsians | 3.51E-04 | 7.38E-04 | 0.634377 |
| *SLC24A4* | rs2402130 | G | A | Europeans | NorthAsians | -2.25E-04 | 7.43E-04 | 0.761965 |
| *OCA2* | rs2311843 | C | T | Europeans | NorthAsians | -0.001944 | 7.40E-04 | **0.008596** |
| *OCA2* | rs1800414 | A | G | Europeans | NorthAsians | -0.002739 | 9.05E-04 | **0.002475** |
| *OCA2* | rs1800404 | C | T | Europeans | NorthAsians | -2.52E-04 | 7.43E-04 | 0.73489 |
| *OCA2* | rs1800401 | C | A | Europeans | NorthAsians | 2.58E-04 | 7.81E-04 | 0.741106 |
| *HERC2* | rs1129038 | G | A | Europeans | NorthAsians | 0.003579 | 8.33E-04 | **1.80E-05** |
| *HERC2* | rs12913832 | A | G | Europeans | NorthAsians | 0.003566 | 8.33E-04 | **1.90E-05** |
| *HERC2* | rs916977 | A | G | Europeans | NorthAsians | 0.001695 | 7.39E-04 | **0.02177** |
| *HERC2* | rs1667394 | G | A | Europeans | NorthAsians | 0.001684 | 7.39E-04 | **0.022651** |
| *SLC24A5* | rs1426654 | G | A | Europeans | NorthAsians | 0.004135 | 7.65E-04 | **0** |
| *MYO5A* | rs4776053 | C | T | Europeans | NorthAsians | -7.63E-04 | 7.38E-04 | 0.301197 |
| *MC1R* | rs2228479 | G | A | Europeans | NorthAsians | -9.75E-04 | 7.40E-04 | 0.187511 |
| *MC1R* | rs885479 | G | A | Europeans | NorthAsians | -0.001829 | 7.39E-04 | **0.013324** |
| *MFSD12* | rs56203814 | C | T | Europeans | NorthAsians | 2.40E-04 | 0.001185 | 0.839641 |
| *MFSD12* | rs10424065 | C | T | Europeans | NorthAsians | 2.40E-04 | 0.001185 | 0.839641 |
| *MFSD12* | rs6510760 | G | A | Europeans | NorthAsians | 1.39E-04 | 7.58E-04 | 0.85424 |
| *MFSD12* | rs112332856 | T | C | Europeans | NorthAsians | 9.48E-04 | 8.98E-04 | 0.291189 |
| *ASIP* | rs6119471 | G | C | Europeans | NorthAsians | 0.001135 | 0.001449 | 0.433361 |
| *ASIP* | rs6058017 | G | A | Europeans | NorthAsians | -0.001192 | 7.91E-04 | 0.132079 |
| *PIGU* | rs2378249 | A | G | Europeans | NorthAsians | -5.74E-04 | 7.39E-04 | 0.437671 |
| *SLC45A2* | rs16891982 | C | G | Europeans | Oceanians | 0.002754 | 6.98E-04 | **8.00E-05** |
| *SLC45A2* | rs28777 | C | A | Europeans | Oceanians | 0.001774 | 6.71E-04 | **0.008209** |
| *SLC45A2* | rs26722 | C | T | Europeans | Oceanians | -0.001337 | 6.78E-04 | **0.048735** |
| *EXOC2* | rs4959270 | C | A | Europeans | Oceanians | -0.001082 | 6.83E-04 | 0.113007 |
| *TYRP1* | rs1408799 | T | C | Europeans | Oceanians | 0.002211 | 7.75E-04 | **0.00434** |
| *TYRP1* | rs683 | A | C | Europeans | Oceanians | -0.002211 | 7.75E-04 | **0.00434** |
| *BNC2* | rs10756819 | G | A | Europeans | Oceanians | 3.18E-04 | 6.68E-04 | 0.633414 |
| *BNC2* | rs12350739 | G | A | Europeans | Oceanians | 0.002508 | 9.67E-04 | **0.009475** |
| *DDB1* | rs11230664 | C | T | Europeans | Oceanians | 0.002313 | 6.83E-04 | **7.03E-04** |
| *DDB1* | rs7120594 | T | C | Europeans | Oceanians | 0.002585 | 6.94E-04 | **1.94E-04** |
| *DDB1* | rs7948623 | A | T | Europeans | Oceanians | -0.002567 | 6.98E-04 | **2.36E-04** |
| *DDB1* | rs1377457 | C | A | Europeans | Oceanians | 0.002405 | 6.86E-04 | **4.53E-04** |
| *TPCN2* | rs35264875 | A | T | Europeans | Oceanians | -5.25E-04 | 6.68E-04 | 0.431752 |
| *TPCN2* | rs3829241 | G | A | Europeans | Oceanians | 0.001206 | 7.11E-04 | 0.090003 |
| *TYR* | rs1042602 | C | A | Europeans | Oceanians | 0.002074 | 9.67E-04 | **0.031884** |
| *TYR* | rs1393350 | G | A | Europeans | Oceanians | 0.001756 | 9.67E-04 | 0.069267 |
| *TYR* | rs1126809 | G | A | Europeans | Oceanians | 0.00178 | 9.67E-04 | 0.065616 |
| *KITLG* | rs642742 | A | G | Europeans | Oceanians | 1.05E-04 | 6.74E-04 | 0.876062 |
| *KITLG* | rs12821256 | T | C | Europeans | Oceanians | 0.001487 | 9.67E-04 | 0.124028 |
| *DCT* | rs1407995 | T | C | Europeans | Oceanians | 6.48E-04 | 6.68E-04 | 0.332199 |
| *DCT* | rs2031526 | G | A | Europeans | Oceanians | -6.46E-04 | 6.68E-04 | 0.334037 |
| *SLC24A4* | rs12896399 | G | T | Europeans | Oceanians | -2.53E-04 | 6.68E-04 | 0.704428 |
| *SLC24A4* | rs2402130 | G | A | Europeans | Oceanians | -9.90E-05 | 6.78E-04 | 0.884527 |
| *OCA2* | rs2311843 | C | T | Europeans | Oceanians | -0.001052 | 6.68E-04 | 0.115666 |
| *OCA2* | rs1800414 | A | G | Europeans | Oceanians | -0.002783 | 7.87E-04 | **4.05E-04** |
| *OCA2* | rs1800404 | C | T | Europeans | Oceanians | 0.001137 | 6.70E-04 | 0.08982 |
| *OCA2* | rs1800401 | C | T | Europeans | Oceanians | 8.27E-04 | 9.69E-04 | 0.393278 |
| *HERC2* | rs1129038 | G | A | Europeans | Oceanians | 0.003037 | 9.67E-04 | **0.001678** |
| *HERC2* | rs12913832 | A | G | Europeans | Oceanians | 0.003027 | 9.67E-04 | **0.001739** |
| *HERC2* | rs916977 | A | G | Europeans | Oceanians | 9.80E-04 | 6.68E-04 | 0.142565 |
| *HERC2* | rs1667394 | G | A | Europeans | Oceanians | 9.71E-04 | 6.68E-04 | 0.14624 |
| *SLC24A5* | rs1426654 | G | A | Europeans | Oceanians | 0.004289 | 7.89E-04 | **0** |
| *MYO5A* | rs4776053 | C | T | Europeans | Oceanians | 9.60E-04 | 7.76E-04 | 0.215663 |
| *MC1R* | rs2228479 | G | A | Europeans | Oceanians | -1.59E-04 | 6.87E-04 | 0.816976 |
| *MC1R* | rs885479 | G | A | Europeans | Oceanians | 4.12E-04 | 7.32E-04 | 0.57365 |
| *MFSD12* | rs56203814 | C | T | Europeans | Oceanians | -4.43E-04 | 9.95E-04 | 0.655763 |
| *MFSD12* | rs10424065 | C | T | Europeans | Oceanians | -4.43E-04 | 9.95E-04 | 0.655763 |
| *MFSD12* | rs6510760 | G | A | Europeans | Oceanians | -0.001365 | 6.69E-04 | **0.041418** |
| *MFSD12* | rs112332856 | T | C | Europeans | Oceanians | -0.001282 | 6.75E-04 | 0.057377 |
| *ASIP* | rs6119471 | G | C | Europeans | Oceanians | 0.001543 | 0.001197 | 0.197389 |
| *ASIP* | rs6058017 | G | A | Europeans | Oceanians | 1.57E-04 | 6.79E-04 | 0.816854 |
| *PIGU* | rs2378249 | A | G | Europeans | Oceanians | -6.90E-05 | 6.81E-04 | 0.9196 |
| *SLC45A2* | rs16891982 | C | G | Europeans | EastAsians | 0.004804 | 7.53E-04 | **0** |
| *SLC45A2* | rs28777 | C | A | Europeans | EastAsians | 0.003301 | 7.40E-04 | **8.00E-06** |
| *SLC45A2* | rs26722 | C | T | Europeans | EastAsians | -0.002147 | 7.43E-04 | **0.00388** |
| *EXOC2* | rs4959270 | C | A | Europeans | EastAsians | 2.80E-04 | 7.35E-04 | 0.70332 |
| *TYRP1* | rs1408799 | T | C | Europeans | EastAsians | 0.002875 | 7.42E-04 | **1.06E-04** |
| *TYRP1* | rs683 | A | C | Europeans | EastAsians | -0.003298 | 7.48E-04 | **1.10E-05** |
| *BNC2* | rs10756819 | G | A | Europeans | EastAsians | 2.60E-04 | 7.35E-04 | 0.723261 |
| *BNC2* | rs12350739 | G | A | Europeans | EastAsians | 0.003316 | 7.58E-04 | **1.20E-05** |
| *DDB1* | rs11230664 | C | T | Europeans | EastAsians | 8.84E-04 | 7.60E-04 | 0.244405 |
| *DDB1* | rs7120594 | T | C | Europeans | EastAsians | 0.001224 | 7.75E-04 | 0.114356 |
| *DDB1* | rs7948623 | A | T | Europeans | EastAsians | 5.27E-04 | 8.46E-04 | 0.533572 |
| *DDB1* | rs1377457 | C | A | Europeans | EastAsians | -2.40E-05 | 7.80E-04 | 0.97525 |
| *TPCN2* | rs35264875 | A | T | Europeans | EastAsians | 0.002125 | 7.51E-04 | **0.004686** |
| *TPCN2* | rs3829241 | G | A | Europeans | EastAsians | 5.75E-04 | 7.35E-04 | 0.43454 |
| *TYR* | rs1042602 | C | A | Europeans | EastAsians | 0.003703 | 8.34E-04 | **9.00E-06** |
| *TYR* | rs1393350 | G | A | Europeans | EastAsians | 0.003095 | 8.07E-04 | **1.26E-04** |
| *TYR* | rs1126809 | G | A | Europeans | EastAsians | 0.003655 | 8.95E-04 | **4.40E-05** |
| *KITLG* | rs642742 | A | G | Europeans | EastAsians | 1.58E-04 | 7.36E-04 | 0.829732 |
| *KITLG* | rs12821256 | T | C | Europeans | EastAsians | 0.003289 | 8.95E-04 | **2.38E-04** |
| *DCT* | rs1407995 | T | C | Europeans | EastAsians | 0.001625 | 7.36E-04 | **0.027173** |
| *DCT* | rs2031526 | G | A | Europeans | EastAsians | -0.001627 | 7.36E-04 | **0.026976** |
| *SLC24A4* | rs12896399 | G | T | Europeans | EastAsians | 3.12E-04 | 7.35E-04 | 0.67082 |
| *SLC24A4* | rs2402130 | G | A | Europeans | EastAsians | -7.69E-04 | 7.37E-04 | 0.296629 |
| *OCA2* | rs2311843 | C | T | Europeans | EastAsians | -0.00198 | 7.36E-04 | **0.007154** |
| *OCA2* | rs1800414 | A | G | Europeans | EastAsians | -0.004515 | 8.94E-04 | **0** |
| *OCA2* | rs1800404 | C | T | Europeans | EastAsians | 0.001211 | 7.35E-04 | 0.099601 |
| *OCA2* | rs1800401 | C | T | Europeans | EastAsians | 0.001064 | 7.57E-04 | 0.160263 |
| *HERC2* | rs1129038 | G | A | Europeans | EastAsians | 0.004539 | 7.92E-04 | **0** |
| *HERC2* | rs12913832 | A | G | Europeans | EastAsians | 0.004684 | 8.07E-04 | **0** |
| *HERC2* | rs916977 | A | G | Europeans | EastAsians | 0.001788 | 7.36E-04 | **0.015136** |
| *HERC2* | rs1667394 | G | A | Europeans | EastAsians | 0.001778 | 7.36E-04 | **0.015681** |
| *SLC24A5* | rs1426654 | G | A | Europeans | EastAsians | 0.005774 | 7.69E-04 | **0** |
| *MYO5A* | rs4776053 | C | T | Europeans | EastAsians | -1.45E-04 | 7.36E-04 | 0.844254 |
| *MC1R* | rs2228479 | G | A | Europeans | EastAsians | -9.51E-04 | 7.37E-04 | 0.196783 |
| *MC1R* | rs885479 | G | A | Europeans | EastAsians | -0.001785 | 7.37E-04 | **0.015392** |
| *MFSD12* | rs56203814 | C | T | Europeans | EastAsians | 0.001563 | 0.001186 | 0.18755 |
| *MFSD12* | rs10424065 | C | T | Europeans | EastAsians | 8.76E-04 | 9.41E-04 | 0.351953 |
| *MFSD12* | rs6510760 | G | A | Europeans | EastAsians | 1.37E-04 | 7.40E-04 | 0.85293 |
| *MFSD12* | rs112332856 | T | C | Europeans | EastAsians | 0.001151 | 7.69E-04 | 0.134437 |
| *ASIP* | rs6119471 | G | C | Europeans | EastAsians | -1.88E-04 | 0.00145 | 0.896809 |
| *ASIP* | rs6058017 | G | A | Europeans | EastAsians | 4.96E-04 | 7.36E-04 | 0.500451 |
| *PIGU* | rs2378249 | A | G | Europeans | EastAsians | -2.14E-04 | 7.36E-04 | 0.770841 |
| *SLC45A2* | rs16891982 | C | G | Europeans | EastAfricans | 0.002845 | 6.31E-04 | **6.00E-06** |
| *SLC45A2* | rs28777 | C | A | Europeans | EastAfricans | 0.001699 | 5.53E-04 | **0.002098** |
| *SLC45A2* | rs26722 | C | T | Europeans | EastAfricans | -4.21E-04 | 5.68E-04 | 0.459218 |
| *EXOC2* | rs4959270 | C | A | Europeans | EastAfricans | 9.79E-04 | 5.64E-04 | 0.08264 |
| *TYRP1* | rs1408799 | T | C | Europeans | EastAfricans | 7.12E-04 | 5.50E-04 | 0.195475 |
| *TYRP1* | rs683 | A | C | Europeans | EastAfricans | -8.09E-04 | 5.51E-04 | 0.142307 |
| *BNC2* | rs10756819 | G | A | Europeans | EastAfricans | 0.001721 | 5.97E-04 | **0.003941** |
| *BNC2* | rs12350739 | G | A | Europeans | EastAfricans | 0.002164 | 7.70E-04 | **0.004947** |
| *DDB1* | rs11230664 | C | T | Europeans | EastAfricans | 0.002891 | 5.75E-04 | **0** |
| *DDB1* | rs7120594 | T | C | Europeans | EastAfricans | 0.002582 | 5.70E-04 | **6.00E-06** |
| *DDB1* | rs7948623 | A | T | Europeans | EastAfricans | -0.002338 | 5.71E-04 | **4.20E-05** |
| *DDB1* | rs1377457 | C | A | Europeans | EastAfricans | 0.003024 | 5.80E-04 | **0** |
| *TPCN2* | rs35264875 | A | T | Europeans | EastAfricans | 8.60E-05 | 5.52E-04 | 0.876402 |
| *TPCN2* | rs3829241 | G | A | Europeans | EastAfricans | 0.001304 | 5.97E-04 | **0.028911** |
| *TYR* | rs1042602 | C | A | Europeans | EastAfricans | 0.001831 | 7.70E-04 | **0.017436** |
| *TYR* | rs1393350 | G | A | Europeans | EastAfricans | 0.001586 | 7.70E-04 | **0.039444** |
| *TYR* | rs1126809 | G | A | Europeans | EastAfricans | 0.001604 | 7.70E-04 | **0.037246** |
| *KITLG* | rs642742 | A | G | Europeans | EastAfricans | 0.001378 | 5.55E-04 | **0.013057** |
| *KITLG* | rs12821256 | T | C | Europeans | EastAfricans | 2.86E-04 | 5.62E-04 | 0.611079 |
| *DCT* | rs1407995 | T | C | Europeans | EastAfricans | -4.00E-05 | 5.53E-04 | 0.942694 |
| *DCT* | rs2031526 | G | A | Europeans | EastAfricans | 6.06E-04 | 5.72E-04 | 0.289335 |
| *SLC24A4* | rs12896399 | G | T | Europeans | EastAfricans | 9.89E-04 | 5.64E-04 | 0.079557 |
| *SLC24A4* | rs2402130 | G | A | Europeans | EastAfricans | 5.86E-04 | 5.49E-04 | 0.285706 |
| *OCA2* | rs2311843 | C | T | Europeans | EastAfricans | 9.39E-04 | 6.29E-04 | 0.135776 |
| *OCA2* | rs1800414 | A | G | Europeans | EastAfricans | -5.29E-04 | 8.31E-04 | 0.524534 |
| *OCA2* | rs1800404 | C | T | Europeans | EastAfricans | 0.001814 | 5.75E-04 | **0.001596** |
| *OCA2* | rs1800401 | C | T | Europeans | EastAfricans | 1.30E-05 | 5.77E-04 | 0.982 |
| *HERC2* | rs1129038 | G | A | Europeans | EastAfricans | 0.002572 | 7.70E-04 | **8.41E-04** |
| *HERC2* | rs12913832 | A | G | Europeans | EastAfricans | 0.002564 | 7.70E-04 | **8.72E-04** |
| *HERC2* | rs916977 | A | G | Europeans | EastAfricans | 0.002182 | 5.97E-04 | **2.60E-04** |
| *HERC2* | rs1667394 | G | A | Europeans | EastAfricans | 0.002175 | 5.97E-04 | **2.72E-04** |
| *SLC24A5* | rs1426654 | G | A | Europeans | EastAfricans | 0.003205 | 5.94E-04 | **0** |
| *MYO5A* | rs4776053 | C | T | Europeans | EastAfricans | 3.64E-04 | 5.65E-04 | 0.518997 |
| *MC1R* | rs2228479 | G | A | Europeans | EastAfricans | 0.001173 | 7.71E-04 | 0.128136 |
| *MC1R* | rs885479 | G | A | Europeans | EastAfricans | 0.001183 | 7.71E-04 | 0.124649 |
| *MFSD12* | rs56203814 | C | T | Europeans | EastAfricans | -0.001548 | 5.81E-04 | **0.007776** |
| *MFSD12* | rs10424065 | C | T | Europeans | EastAfricans | -0.002352 | 5.78E-04 | **4.60E-05** |
| *MFSD12* | rs6510760 | G | A | Europeans | EastAfricans | -0.002269 | 5.76E-04 | **8.10E-05** |
| *MFSD12* | rs112332856 | T | C | Europeans | EastAfricans | -0.00259 | 5.77E-04 | **7.00E-06** |
| *ASIP* | rs6119471 | G | C | Europeans | EastAfricans | 0.002737 | 7.73E-04 | **3.98E-04** |
| *ASIP* | rs6058017 | G | A | Europeans | EastAfricans | 0.001219 | 5.51E-04 | **0.026928** |
| *PIGU* | rs2378249 | A | G | Europeans | EastAfricans | 3.14E-04 | 5.65E-04 | 0.578311 |
| *SLC45A2* | rs16891982 | C | G | Europeans | WestAfricans | 0.00263 | 4.79E-04 | **0** |
| *SLC45A2* | rs28777 | C | A | Europeans | WestAfricans | 0.001508 | 4.52E-04 | **8.40E-04** |
| *SLC45A2* | rs26722 | C | T | Europeans | WestAfricans | -2.48E-04 | 4.55E-04 | 0.585986 |
| *EXOC2* | rs4959270 | C | A | Europeans | WestAfricans | 3.28E-04 | 4.50E-04 | 0.465309 |
| *TYRP1* | rs1408799 | T | C | Europeans | WestAfricans | 6.83E-04 | 4.50E-04 | 0.128978 |
| *TYRP1* | rs683 | A | C | Europeans | WestAfricans | -8.97E-04 | 4.50E-04 | **0.046465** |
| *BNC2* | rs10756819 | G | A | Europeans | WestAfricans | 9.96E-04 | 4.51E-04 | **0.027057** |
| *BNC2* | rs12350739 | G | A | Europeans | WestAfricans | 0.00186 | 4.88E-04 | **1.39E-04** |
| *DDB1* | rs11230664 | C | T | Europeans | WestAfricans | 0.001682 | 4.58E-04 | **2.38E-04** |
| *DDB1* | rs7120594 | T | C | Europeans | WestAfricans | 0.001719 | 4.64E-04 | **2.09E-04** |
| *DDB1* | rs7948623 | A | T | Europeans | WestAfricans | -0.001286 | 4.66E-04 | **0.005809** |
| *DDB1* | rs1377457 | C | A | Europeans | WestAfricans | 0.001757 | 4.59E-04 | **1.31E-04** |
| *TPCN2* | rs35264875 | A | T | Europeans | WestAfricans | 0.001083 | 4.64E-04 | **0.019467** |
| *TPCN2* | rs3829241 | G | A | Europeans | WestAfricans | 9.58E-04 | 4.53E-04 | 0.034415 |
| *TYR* | rs1042602 | C | A | Europeans | WestAfricans | 0.001495 | 4.77E-04 | **0.001739** |
| *TYR* | rs1393350 | G | A | Europeans | WestAfricans | 0.001893 | 6.21E-04 | **0.002296** |
| *TYR* | rs1126809 | G | A | Europeans | WestAfricans | 0.001317 | 4.78E-04 | **0.005828** |
| *KITLG* | rs642742 | A | G | Europeans | WestAfricans | 0.001239 | 4.51E-04 | **0.005976** |
| *KITLG* | rs12821256 | T | C | Europeans | WestAfricans | 0.00173 | 6.21E-04 | **0.005338** |
| *DCT* | rs1407995 | T | C | Europeans | WestAfricans | 6.70E-05 | 4.50E-04 | 0.881067 |
| *DCT* | rs2031526 | G | A | Europeans | WestAfricans | 3.96E-04 | 4.51E-04 | 0.380466 |
| *SLC24A4* | rs12896399 | G | T | Europeans | WestAfricans | 0.001753 | 4.88E-04 | **3.32E-04** |
| *SLC24A4* | rs2402130 | G | A | Europeans | WestAfricans | 5.91E-04 | 4.50E-04 | 0.188372 |
| *OCA2* | rs2311843 | C | T | Europeans | WestAfricans | 5.25E-04 | 4.54E-04 | 0.246673 |
| *OCA2* | rs1800414 | A | G | Europeans | WestAfricans | 2.26E-04 | 6.68E-04 | 0.734638 |
| *OCA2* | rs1800404 | C | T | Europeans | WestAfricans | 0.001079 | 4.50E-04 | **0.016588** |
| *OCA2* | rs1800401 | C | T | Europeans | WestAfricans | -4.49E-04 | 4.51E-04 | 0.320382 |
| *HERC2* | rs1129038 | G | A | Europeans | WestAfricans | 0.00197 | 4.69E-04 | **2.70E-05** |
| *HERC2* | rs12913832 | A | G | Europeans | WestAfricans | 0.001964 | 4.69E-04 | **2.80E-05** |
| *HERC2* | rs916977 | A | G | Europeans | WestAfricans | 0.001317 | 4.51E-04 | **0.003482** |
| *HERC2* | rs1667394 | G | A | Europeans | WestAfricans | 0.001354 | 4.51E-04 | **0.002681** |
| *SLC24A5* | rs1426654 | G | A | Europeans | WestAfricans | 0.002367 | 4.60E-04 | **0** |
| *MYO5A* | rs4776053 | C | T | Europeans | WestAfricans | 1.97E-04 | 4.51E-04 | 0.661346 |
| *MC1R* | rs2228479 | G | A | Europeans | WestAfricans | 0.001567 | 6.21E-04 | **0.011649** |
| *MC1R* | rs885479 | G | A | Europeans | WestAfricans | 0.001576 | 6.21E-04 | **0.011201** |
| *MFSD12* | rs56203814 | C | T | Europeans | WestAfricans | -0.001347 | 4.72E-04 | **0.004295** |
| *MFSD12* | rs10424065 | C | T | Europeans | WestAfricans | -0.001503 | 4.71E-04 | **0.001434** |
| *MFSD12* | rs6510760 | G | A | Europeans | WestAfricans | -0.001407 | 4.51E-04 | **0.001801** |
| *MFSD12* | rs112332856 | T | C | Europeans | WestAfricans | -0.001501 | 4.52E-04 | **8.93E-04** |
| *ASIP* | rs6119471 | G | C | Europeans | WestAfricans | 0.002735 | 6.21E-04 | **1.10E-05** |
| *ASIP* | rs6058017 | G | A | Europeans | WestAfricans | 0.001095 | 4.50E-04 | **0.015015** |
| *PIGU* | rs2378249 | A | G | Europeans | WestAfricans | -8.40E-05 | 4.50E-04 | 0.852618 |
| *SLC45A2* | rs16891982 | C | G | NorthAsians | Oceanians | 2.09E-04 | 6.90E-04 | 0.761863 |
| *SLC45A2* | rs28777 | C | A | NorthAsians | Oceanians | -7.11E-04 | 6.59E-04 | 0.280548 |
| *SLC45A2* | rs26722 | C | T | NorthAsians | Oceanians | 7.70E-05 | 6.62E-04 | 0.907271 |
| *EXOC2* | rs4959270 | C | A | NorthAsians | Oceanians | -0.001411 | 6.72E-04 | **0.03568** |
| *TYRP1* | rs1408799 | T | C | NorthAsians | Oceanians | -8.50E-05 | 7.99E-04 | 0.915284 |
| *TYRP1* | rs683 | A | C | NorthAsians | Oceanians | -3.28E-04 | 7.79E-04 | 0.67392 |
| *BNC2* | rs10756819 | G | A | NorthAsians | Oceanians | 1.32E-04 | 6.56E-04 | 0.840146 |
| *BNC2* | rs12350739 | G | A | NorthAsians | Oceanians | 5.27E-04 | 9.83E-04 | 0.591614 |
| *DDB1* | rs11230664 | C | T | NorthAsians | Oceanians | 0.002999 | 9.63E-04 | **0.001845** |
| *DDB1* | rs7120594 | T | C | NorthAsians | Oceanians | 0.002999 | 9.63E-04 | **0.001845** |
| *DDB1* | rs7948623 | A | T | NorthAsians | Oceanians | -0.002904 | 9.63E-04 | **0.00257** |
| *DDB1* | rs1377457 | C | A | NorthAsians | Oceanians | 0.002999 | 9.63E-04 | **0.001845** |
| *TPCN2* | rs35264875 | A | T | NorthAsians | Oceanians | -0.001741 | 6.91E-04 | **0.011742** |
| *TPCN2* | rs3829241 | G | A | NorthAsians | Oceanians | 0.001063 | 7.01E-04 | 0.12938 |
| *TYR* | rs1042602 | C | A | NorthAsians | Oceanians | 4.72E-04 | 9.85E-04 | 0.631774 |
| *TYR* | rs1393350 | G | A | NorthAsians | Oceanians | -8.40E-05 | 0.00104 | 0.935849 |
| *TYR* | rs1126809 | G | A | NorthAsians | Oceanians | -8.40E-05 | 0.00104 | 0.935849 |
| *KITLG* | rs642742 | A | G | NorthAsians | Oceanians | 2.60E-04 | 6.66E-04 | 0.696333 |
| *KITLG* | rs12821256 | T | C | NorthAsians | Oceanians | -6.35E-04 | 0.00119 | 0.593386 |
| *DCT* | rs1407995 | T | C | NorthAsians | Oceanians | -5.75E-04 | 6.57E-04 | 0.381764 |
| *DCT* | rs2031526 | G | A | NorthAsians | Oceanians | 5.75E-04 | 6.57E-04 | 0.381764 |
| *SLC24A4* | rs12896399 | G | T | NorthAsians | Oceanians | -5.34E-04 | 6.57E-04 | 0.416225 |
| *SLC24A4* | rs2402130 | G | A | NorthAsians | Oceanians | 8.10E-05 | 6.70E-04 | 0.903342 |
| *OCA2* | rs2311843 | C | T | NorthAsians | Oceanians | 5.03E-04 | 6.58E-04 | 0.444072 |
| *OCA2* | rs1800414 | A | G | NorthAsians | Oceanians | -5.92E-04 | 6.70E-04 | 0.376725 |
| *OCA2* | rs1800404 | C | T | NorthAsians | Oceanians | 0.001338 | 6.62E-04 | **0.04329** |
| *OCA2* | rs1800401 | C | T | NorthAsians | Oceanians | 6.21E-04 | 9.78E-04 | 0.525882 |
| *HERC2* | rs1129038 | G | A | NorthAsians | Oceanians | 1.74E-04 | 0.001008 | 0.862992 |
| *HERC2* | rs12913832 | A | G | NorthAsians | Oceanians | 1.74E-04 | 0.001008 | 0.862992 |
| *HERC2* | rs916977 | A | G | NorthAsians | Oceanians | -3.76E-04 | 6.57E-04 | 0.567236 |
| *HERC2* | rs1667394 | G | A | NorthAsians | Oceanians | -3.76E-04 | 6.57E-04 | 0.567236 |
| *SLC24A5* | rs1426654 | G | A | NorthAsians | Oceanians | 9.81E-04 | 7.68E-04 | 0.201703 |
| *MYO5A* | rs4776053 | C | T | NorthAsians | Oceanians | 0.001571 | 7.65E-04 | **0.04014** |
| *MC1R* | rs2228479 | G | A | NorthAsians | Oceanians | 6.21E-04 | 6.75E-04 | 0.357846 |
| *MC1R* | rs885479 | G | A | NorthAsians | Oceanians | 0.001875 | 7.21E-04 | **0.009264** |
| *MFSD12* | rs56203814 | C | T | NorthAsians | Oceanians | -6.35E-04 | 0.00119 | 0.593386 |
| *MFSD12* | rs10424065 | C | T | NorthAsians | Oceanians | -6.35E-04 | 0.00119 | 0.593386 |
| *MFSD12* | rs6510760 | G | A | NorthAsians | Oceanians | -0.001476 | 6.69E-04 | **0.027402** |
| *MFSD12* | rs112332856 | T | C | NorthAsians | Oceanians | -0.002041 | 7.73E-04 | **0.008293** |
| *ASIP* | rs6119471 | G | C | NorthAsians | Oceanians | 6.35E-04 | 0.00119 | 0.593386 |
| *ASIP* | rs6058017 | G | A | NorthAsians | Oceanians | 0.001111 | 7.05E-04 | 0.115076 |
| *PIGU* | rs2378249 | A | G | NorthAsians | Oceanians | 3.90E-04 | 6.70E-04 | 0.560041 |
| *SLC45A2* | rs16891982 | C | G | NorthAsians | EastAsians | 0.001997 | 5.71E-04 | **4.71E-04** |
| *SLC45A2* | rs28777 | C | A | NorthAsians | EastAsians | 2.38E-04 | 5.38E-04 | 0.657895 |
| *SLC45A2* | rs26722 | C | T | NorthAsians | EastAsians | -4.67E-04 | 5.31E-04 | 0.378937 |
| *EXOC2* | rs4959270 | C | A | NorthAsians | EastAsians | -1.62E-04 | 5.30E-04 | 0.759375 |
| *TYRP1* | rs1408799 | T | C | NorthAsians | EastAsians | 6.00E-06 | 6.46E-04 | 0.992184 |
| *TYRP1* | rs683 | A | C | NorthAsians | EastAsians | -0.001162 | 6.01E-04 | 0.053093 |
| *BNC2* | rs10756819 | G | A | NorthAsians | EastAsians | 3.40E-05 | 5.28E-04 | 0.948733 |
| *BNC2* | rs12350739 | G | A | NorthAsians | EastAsians | 0.001035 | 6.65E-04 | 0.119833 |
| *DDB1* | rs11230664 | C | T | NorthAsians | EastAsians | 0.002143 | 0.001209 | 0.076471 |
| *DDB1* | rs7120594 | T | C | NorthAsians | EastAsians | 0.002143 | 0.001209 | 0.076471 |
| *DDB1* | rs7948623 | A | T | NorthAsians | EastAsians | 1.30E-04 | 0.001274 | 0.918425 |
| *DDB1* | rs1377457 | C | A | NorthAsians | EastAsians | 8.83E-04 | 0.001224 | 0.470813 |
| *TPCN2* | rs35264875 | A | T | NorthAsians | EastAsians | 7.44E-04 | 6.52E-04 | 0.253889 |
| *TPCN2* | rs3829241 | G | A | NorthAsians | EastAsians | 4.87E-04 | 5.30E-04 | 0.358337 |
| *TYR* | rs1042602 | C | A | NorthAsians | EastAsians | 0.002093 | 7.99E-04 | **0.008792** |
| *TYR* | rs1393350 | G | A | NorthAsians | EastAsians | 9.79E-04 | 9.13E-04 | 0.28368 |
| *TYR* | rs1126809 | G | A | NorthAsians | EastAsians | 0.001632 | 0.001029 | 0.112903 |
| *KITLG* | rs642742 | A | G | NorthAsians | EastAsians | 4.33E-04 | 5.40E-04 | 0.422821 |
| *KITLG* | rs12821256 | T | C | NorthAsians | EastAsians | 7.83E-04 | 0.001359 | 0.564616 |
| *DCT* | rs1407995 | T | C | NorthAsians | EastAsians | 1.19E-04 | 5.31E-04 | 0.822512 |
| *DCT* | rs2031526 | G | A | NorthAsians | EastAsians | -1.25E-04 | 5.31E-04 | 0.813192 |
| *SLC24A4* | rs12896399 | G | T | NorthAsians | EastAsians | -4.70E-05 | 5.30E-04 | 0.928955 |
| *SLC24A4* | rs2402130 | G | A | NorthAsians | EastAsians | -6.70E-04 | 5.43E-04 | 0.217046 |
| *OCA2* | rs2311843 | C | T | NorthAsians | EastAsians | -4.50E-05 | 5.33E-04 | 0.933321 |
| *OCA2* | rs1800414 | A | G | NorthAsians | EastAsians | -0.002186 | 5.52E-04 | **7.40E-05** |
| *OCA2* | rs1800404 | C | T | NorthAsians | EastAsians | 0.0018 | 5.40E-04 | **8.52E-04** |
| *OCA2* | rs1800401 | C | T | NorthAsians | EastAsians | 9.92E-04 | 6.41E-04 | 0.122014 |
| *HERC2* | rs1129038 | G | A | NorthAsians | EastAsians | 0.001182 | 7.99E-04 | 0.139204 |
| *HERC2* | rs12913832 | A | G | NorthAsians | EastAsians | 0.001375 | 8.22E-04 | 0.094386 |
| *HERC2* | rs916977 | A | G | NorthAsians | EastAsians | 1.14E-04 | 5.30E-04 | 0.829383 |
| *HERC2* | rs1667394 | G | A | NorthAsians | EastAsians | 1.16E-04 | 5.30E-04 | 0.826312 |
| *SLC24A5* | rs1426654 | G | A | NorthAsians | EastAsians | 0.002017 | 5.61E-04 | **3.21E-04** |
| *MYO5A* | rs4776053 | C | T | NorthAsians | EastAsians | 7.61E-04 | 5.30E-04 | 0.1509 |
| *MC1R* | rs2228479 | G | A | NorthAsians | EastAsians | 2.90E-05 | 5.30E-04 | 0.956249 |
| *MC1R* | rs885479 | G | A | NorthAsians | EastAsians | 5.40E-05 | 5.29E-04 | 0.918692 |
| *MFSD12* | rs56203814 | C | T | NorthAsians | EastAsians | 0.001629 | 0.001624 | 0.315933 |
| *MFSD12* | rs10424065 | C | T | NorthAsians | EastAsians | 7.83E-04 | 0.001359 | 0.564616 |
| *MFSD12* | rs6510760 | G | A | NorthAsians | EastAsians | -3.00E-06 | 5.71E-04 | 0.996409 |
| *MFSD12* | rs112332856 | T | C | NorthAsians | EastAsians | 2.50E-04 | 8.55E-04 | 0.769913 |
| *ASIP* | rs6119471 | G | C | NorthAsians | EastAsians | -0.001629 | 0.001624 | 0.315933 |
| *ASIP* | rs6058017 | G | A | NorthAsians | EastAsians | 0.002077 | 6.35E-04 | **0.001071** |
| *PIGU* | rs2378249 | A | G | NorthAsians | EastAsians | 4.42E-04 | 5.32E-04 | 0.405868 |
| *SLC45A2* | rs16891982 | C | G | NorthAsians | EastAfricans | 8.87E-04 | 7.11E-04 | 0.21234 |
| *SLC45A2* | rs28777 | C | A | NorthAsians | EastAfricans | -2.13E-04 | 6.41E-04 | 0.740327 |
| *SLC45A2* | rs26722 | C | T | NorthAsians | EastAfricans | 6.67E-04 | 6.52E-04 | 0.306577 |
| *EXOC2* | rs4959270 | C | A | NorthAsians | EastAfricans | 7.26E-04 | 6.52E-04 | 0.26576 |
| *TYRP1* | rs1408799 | T | C | NorthAsians | EastAfricans | -0.001054 | 6.63E-04 | 0.11218 |
| *TYRP1* | rs683 | A | C | NorthAsians | EastAfricans | 6.40E-04 | 6.50E-04 | 0.325313 |
| *BNC2* | rs10756819 | G | A | NorthAsians | EastAfricans | 0.001578 | 6.80E-04 | **0.020383** |
| *BNC2* | rs12350739 | G | A | NorthAsians | EastAfricans | 6.41E-04 | 8.53E-04 | 0.45233 |
| *DDB1* | rs11230664 | C | T | NorthAsians | EastAfricans | 0.003418 | 8.48E-04 | **5.50E-05** |
| *DDB1* | rs7120594 | T | C | NorthAsians | EastAfricans | 0.0029 | 8.39E-04 | **5.48E-04** |
| *DDB1* | rs7948623 | A | T | NorthAsians | EastAfricans | -0.002597 | 8.38E-04 | **0.001932** |
| *DDB1* | rs1377457 | C | A | NorthAsians | EastAfricans | 0.00348 | 8.50E-04 | **4.20E-05** |
| *TPCN2* | rs35264875 | A | T | NorthAsians | EastAfricans | -8.49E-04 | 6.62E-04 | 0.199751 |
| *TPCN2* | rs3829241 | G | A | NorthAsians | EastAfricans | 0.001194 | 6.80E-04 | 0.079327 |
| *TYR* | rs1042602 | C | A | NorthAsians | EastAfricans | 5.99E-04 | 8.55E-04 | 0.48367 |
| *TYR* | rs1393350 | G | A | NorthAsians | EastAfricans | 1.71E-04 | 8.92E-04 | 0.848096 |
| *TYR* | rs1126809 | G | A | NorthAsians | EastAfricans | 1.71E-04 | 8.92E-04 | 0.848096 |
| *KITLG* | rs642742 | A | G | NorthAsians | EastAfricans | 0.001497 | 6.46E-04 | **0.020482** |
| *KITLG* | rs12821256 | T | C | NorthAsians | EastAfricans | -0.001347 | 8.46E-04 | 0.111491 |
| *DCT* | rs1407995 | T | C | NorthAsians | EastAfricans | -9.80E-04 | 6.42E-04 | 0.126771 |
| *DCT* | rs2031526 | G | A | NorthAsians | EastAfricans | 0.001545 | 6.59E-04 | **0.019011** |
| *SLC24A4* | rs12896399 | G | T | NorthAsians | EastAfricans | 7.73E-04 | 6.52E-04 | 0.235596 |
| *SLC24A4* | rs2402130 | G | A | NorthAsians | EastAfricans | 7.24E-04 | 6.40E-04 | 0.258123 |
| *OCA2* | rs2311843 | C | T | NorthAsians | EastAfricans | 0.002135 | 7.09E-04 | **0.002614** |
| *OCA2* | rs1800414 | A | G | NorthAsians | EastAfricans | 0.001157 | 8.40E-04 | 0.168583 |
| *OCA2* | rs1800404 | C | T | NorthAsians | EastAfricans | 0.001969 | 6.63E-04 | **0.002979** |
| *OCA2* | rs1800401 | C | T | NorthAsians | EastAfricans | -1.46E-04 | 6.78E-04 | 0.829763 |
| *HERC2* | rs1129038 | G | A | NorthAsians | EastAfricans | 3.69E-04 | 8.70E-04 | 0.671346 |
| *HERC2* | rs12913832 | A | G | NorthAsians | EastAfricans | 3.69E-04 | 8.70E-04 | 0.671346 |
| *HERC2* | rs916977 | A | G | NorthAsians | EastAfricans | 0.001139 | 6.81E-04 | 0.094325 |
| *HERC2* | rs1667394 | G | A | NorthAsians | EastAfricans | 0.001139 | 6.81E-04 | 0.094325 |
| *SLC24A5* | rs1426654 | G | A | NorthAsians | EastAfricans | 6.60E-04 | 6.70E-04 | 0.324444 |
| *MYO5A* | rs4776053 | C | T | NorthAsians | EastAfricans | 8.34E-04 | 6.52E-04 | 0.200935 |
| *MC1R* | rs2228479 | G | A | NorthAsians | EastAfricans | 0.001772 | 8.37E-04 | **0.034127** |
| *MC1R* | rs885479 | G | A | NorthAsians | EastAfricans | 0.002309 | 8.36E-04 | **0.005774** |
| *MFSD12* | rs56203814 | C | T | NorthAsians | EastAfricans | -0.001695 | 8.40E-04 | **0.043614** |
| *MFSD12* | rs10424065 | C | T | NorthAsians | EastAfricans | -0.0025 | 8.37E-04 | **0.002835** |
| *MFSD12* | rs6510760 | G | A | NorthAsians | EastAfricans | -0.002355 | 6.68E-04 | **4.27E-04** |
| *MFSD12* | rs112332856 | T | C | NorthAsians | EastAfricans | -0.003174 | 7.30E-04 | **1.40E-05** |
| *ASIP* | rs6119471 | G | C | NorthAsians | EastAfricans | 0.002038 | 8.38E-04 | **0.014996** |
| *ASIP* | rs6058017 | G | A | NorthAsians | EastAfricans | 0.001952 | 6.63E-04 | **0.003258** |
| *PIGU* | rs2378249 | A | G | NorthAsians | EastAfricans | 6.67E-04 | 6.52E-04 | 0.306577 |
| *SLC45A2* | rs16891982 | C | G | NorthAsians | WestAfricans | 0.001087 | 5.44E-04 | **0.045692** |
| *SLC45A2* | rs28777 | C | A | NorthAsians | WestAfricans | 2.00E-06 | 5.19E-04 | 0.997565 |
| *SLC45A2* | rs26722 | C | T | NorthAsians | WestAfricans | 6.09E-04 | 5.20E-04 | 0.240886 |
| *EXOC2* | rs4959270 | C | A | NorthAsians | WestAfricans | 1.29E-04 | 5.18E-04 | 0.803923 |
| *TYRP1* | rs1408799 | T | C | NorthAsians | WestAfricans | -7.09E-04 | 5.36E-04 | 0.185954 |
| *TYRP1* | rs683 | A | C | NorthAsians | WestAfricans | 2.45E-04 | 5.26E-04 | 0.641416 |
| *BNC2* | rs10756819 | G | A | NorthAsians | WestAfricans | 8.84E-04 | 5.19E-04 | 0.088369 |
| *BNC2* | rs12350739 | G | A | NorthAsians | WestAfricans | 6.60E-04 | 5.67E-04 | 0.244361 |
| *DDB1* | rs11230664 | C | T | NorthAsians | WestAfricans | 0.002097 | 6.71E-04 | **0.001773** |
| *DDB1* | rs7120594 | T | C | NorthAsians | WestAfricans | 0.00197 | 6.71E-04 | **0.003326** |
| *DDB1* | rs7948623 | A | T | NorthAsians | WestAfricans | -0.00149 | 6.71E-04 | **0.026434** |
| *DDB1* | rs1377457 | C | A | NorthAsians | WestAfricans | 0.002116 | 6.71E-04 | **0.001611** |
| *TPCN2* | rs35264875 | A | T | NorthAsians | WestAfricans | 3.46E-04 | 5.46E-04 | 0.525583 |
| *TPCN2* | rs3829241 | G | A | NorthAsians | WestAfricans | 8.71E-04 | 5.21E-04 | 0.094207 |
| *TYR* | rs1042602 | C | A | NorthAsians | WestAfricans | 5.24E-04 | 5.59E-04 | 0.348536 |
| *TYR* | rs1393350 | G | A | NorthAsians | WestAfricans | 7.78E-04 | 7.15E-04 | 0.276486 |
| *TYR* | rs1126809 | G | A | NorthAsians | WestAfricans | 1.87E-04 | 5.95E-04 | 0.752689 |
| *KITLG* | rs642742 | A | G | NorthAsians | WestAfricans | 0.001333 | 5.20E-04 | **0.010391** |
| *KITLG* | rs12821256 | T | C | NorthAsians | WestAfricans | 4.44E-04 | 7.96E-04 | 0.577185 |
| *DCT* | rs1407995 | T | C | NorthAsians | WestAfricans | -6.74E-04 | 5.18E-04 | 0.193307 |
| *DCT* | rs2031526 | G | A | NorthAsians | WestAfricans | 0.001135 | 5.19E-04 | **0.028743** |
| *SLC24A4* | rs12896399 | G | T | NorthAsians | WestAfricans | 0.001583 | 5.52E-04 | **0.004124** |
| *SLC24A4* | rs2402130 | G | A | NorthAsians | WestAfricans | 7.00E-04 | 5.19E-04 | 0.177234 |
| *OCA2* | rs2311843 | C | T | NorthAsians | WestAfricans | 0.001468 | 5.21E-04 | **0.004867** |
| *OCA2* | rs1800414 | A | G | NorthAsians | WestAfricans | 0.001555 | 6.75E-04 | **0.021184** |
| *OCA2* | rs1800404 | C | T | NorthAsians | WestAfricans | 0.001201 | 5.20E-04 | **0.020896** |
| *OCA2* | rs1800401 | C | T | NorthAsians | WestAfricans | -5.74E-04 | 5.31E-04 | 0.280179 |
| *HERC2* | rs1129038 | G | A | NorthAsians | WestAfricans | 2.35E-04 | 5.67E-04 | 0.678681 |
| *HERC2* | rs12913832 | A | G | NorthAsians | WestAfricans | 2.35E-04 | 5.67E-04 | 0.678681 |
| *HERC2* | rs916977 | A | G | NorthAsians | WestAfricans | 4.95E-04 | 5.19E-04 | 0.33946 |
| *HERC2* | rs1667394 | G | A | NorthAsians | WestAfricans | 5.38E-04 | 5.19E-04 | 0.30001 |
| *SLC24A5* | rs1426654 | G | A | NorthAsians | WestAfricans | 3.62E-04 | 5.21E-04 | 0.486252 |
| *MYO5A* | rs4776053 | C | T | NorthAsians | WestAfricans | 5.67E-04 | 5.18E-04 | 0.273699 |
| *MC1R* | rs2228479 | G | A | NorthAsians | WestAfricans | 0.00204 | 6.72E-04 | **0.002397** |
| *MC1R* | rs885479 | G | A | NorthAsians | WestAfricans | 0.002462 | 6.72E-04 | **2.46E-04** |
| *MFSD12* | rs56203814 | C | T | NorthAsians | WestAfricans | -0.001463 | 6.71E-04 | **0.029258** |
| *MFSD12* | rs10424065 | C | T | NorthAsians | WestAfricans | -0.001619 | 6.71E-04 | **0.01582** |
| *MFSD12* | rs6510760 | G | A | NorthAsians | WestAfricans | -0.001475 | 5.24E-04 | **0.004891** |
| *MFSD12* | rs112332856 | T | C | NorthAsians | WestAfricans | -0.001961 | 5.73E-04 | **6.16E-04** |
| *ASIP* | rs6119471 | G | C | NorthAsians | WestAfricans | 0.002184 | 6.71E-04 | **0.001134** |
| *ASIP* | rs6058017 | G | A | NorthAsians | WestAfricans | 0.001673 | 5.36E-04 | **0.001802** |
| *PIGU* | rs2378249 | A | G | NorthAsians | WestAfricans | 1.95E-04 | 5.18E-04 | 0.707335 |
| *SLC45A2* | rs16891982 | C | G | Oceanians | EastAsians | 0.001089 | 6.67E-04 | 0.102701 |
| *SLC45A2* | rs28777 | C | A | Oceanians | EastAsians | 8.66E-04 | 6.28E-04 | 0.167747 |
| *SLC45A2* | rs26722 | C | T | Oceanians | EastAsians | -3.81E-04 | 6.32E-04 | 0.546907 |
| *EXOC2* | rs4959270 | C | A | Oceanians | EastAsians | 0.001306 | 6.43E-04 | **0.042308** |
| *TYRP1* | rs1408799 | T | C | Oceanians | EastAsians | 8.90E-05 | 7.45E-04 | 0.904817 |
| *TYRP1* | rs683 | A | C | Oceanians | EastAsians | -4.27E-04 | 7.49E-04 | 0.568553 |
| *BNC2* | rs10756819 | G | A | Oceanians | EastAsians | -1.10E-04 | 6.27E-04 | 0.860449 |
| *BNC2* | rs12350739 | G | A | Oceanians | EastAsians | 1.45E-04 | 9.51E-04 | 0.8785 |
| *DDB1* | rs11230664 | C | T | Oceanians | EastAsians | -0.001606 | 6.30E-04 | **0.010798** |
| *DDB1* | rs7120594 | T | C | Oceanians | EastAsians | -0.001606 | 6.30E-04 | **0.010798** |
| *DDB1* | rs7948623 | A | T | Oceanians | EastAsians | 0.002989 | 6.82E-04 | **1.20E-05** |
| *DDB1* | rs1377457 | C | A | Oceanians | EastAsians | -0.002425 | 6.42E-04 | **1.59E-04** |
| *TPCN2* | rs35264875 | A | T | Oceanians | EastAsians | 0.002225 | 6.40E-04 | **5.03E-04** |
| *TPCN2* | rs3829241 | G | A | Oceanians | EastAsians | -7.46E-04 | 6.73E-04 | 0.267951 |
| *TYR* | rs1042602 | C | A | Oceanians | EastAsians | 8.89E-04 | 9.91E-04 | 0.369733 |
| *TYR* | rs1393350 | G | A | Oceanians | EastAsians | 7.20E-04 | 9.76E-04 | 0.460701 |
| *TYR* | rs1126809 | G | A | Oceanians | EastAsians | 0.001144 | 0.001024 | 0.26368 |
| *KITLG* | rs642742 | A | G | Oceanians | EastAsians | 2.10E-05 | 6.34E-04 | 0.973094 |
| *KITLG* | rs12821256 | T | C | Oceanians | EastAsians | 0.001144 | 0.001024 | 0.26368 |
| *DCT* | rs1407995 | T | C | Oceanians | EastAsians | 6.52E-04 | 6.28E-04 | 0.298811 |
| *DCT* | rs2031526 | G | A | Oceanians | EastAsians | -6.56E-04 | 6.28E-04 | 0.295756 |
| *SLC24A4* | rs12896399 | G | T | Oceanians | EastAsians | 5.03E-04 | 6.28E-04 | 0.422541 |
| *SLC24A4* | rs2402130 | G | A | Oceanians | EastAsians | -5.17E-04 | 6.40E-04 | 0.418887 |
| *OCA2* | rs2311843 | C | T | Oceanians | EastAsians | -5.32E-04 | 6.28E-04 | 0.396381 |
| *OCA2* | rs1800414 | A | G | Oceanians | EastAsians | -8.29E-04 | 6.33E-04 | 0.190345 |
| *OCA2* | rs1800404 | C | T | Oceanians | EastAsians | -1.68E-04 | 6.29E-04 | 0.789951 |
| *OCA2* | rs1800401 | C | T | Oceanians | EastAsians | 2.40E-05 | 9.48E-04 | 0.979833 |
| *HERC2* | rs1129038 | G | A | Oceanians | EastAsians | 5.94E-04 | 9.68E-04 | 0.539336 |
| *HERC2* | rs12913832 | A | G | Oceanians | EastAsians | 7.20E-04 | 9.76E-04 | 0.460701 |
| *HERC2* | rs916977 | A | G | Oceanians | EastAsians | 4.50E-04 | 6.27E-04 | 0.473124 |
| *HERC2* | rs1667394 | G | A | Oceanians | EastAsians | 4.51E-04 | 6.27E-04 | 0.471789 |
| *SLC24A5* | rs1426654 | G | A | Oceanians | EastAsians | 3.31E-04 | 7.48E-04 | 0.658389 |
| *MYO5A* | rs4776053 | C | T | Oceanians | EastAsians | -0.001076 | 7.41E-04 | 0.146351 |
| *MC1R* | rs2228479 | G | A | Oceanians | EastAsians | -6.02E-04 | 6.47E-04 | 0.351911 |
| *MC1R* | rs885479 | G | A | Oceanians | EastAsians | -0.00184 | 6.94E-04 | **0.008036** |
| *MFSD12* | rs56203814 | C | T | Oceanians | EastAsians | 0.001694 | 0.001175 | 0.149534 |
| *MFSD12* | rs10424065 | C | T | Oceanians | EastAsians | 0.001144 | 0.001024 | 0.26368 |
| *MFSD12* | rs6510760 | G | A | Oceanians | EastAsians | 0.001475 | 6.29E-04 | **0.019026** |
| *MFSD12* | rs112332856 | T | C | Oceanians | EastAsians | 0.002203 | 6.52E-04 | **7.21E-04** |
| *ASIP* | rs6119471 | G | C | Oceanians | EastAsians | -0.001694 | 0.001175 | 0.149534 |
| *ASIP* | rs6058017 | G | A | Oceanians | EastAsians | 2.40E-04 | 6.38E-04 | 0.707433 |
| *PIGU* | rs2378249 | A | G | Oceanians | EastAsians | -1.03E-04 | 6.41E-04 | 0.872447 |
| *SLC45A2* | rs16891982 | C | G | Oceanians | EastAfricans | 7.26E-04 | 7.29E-04 | 0.318758 |
| *SLC45A2* | rs28777 | C | A | Oceanians | EastAfricans | 3.35E-04 | 6.44E-04 | 0.603448 |
| *SLC45A2* | rs26722 | C | T | Oceanians | EastAfricans | 6.08E-04 | 6.59E-04 | 0.356442 |
| *EXOC2* | rs4959270 | C | A | Oceanians | EastAfricans | 0.001811 | 6.65E-04 | **0.006465** |
| *TYRP1* | rs1408799 | T | C | Oceanians | EastAfricans | -9.88E-04 | 7.12E-04 | 0.164947 |
| *TYRP1* | rs683 | A | C | Oceanians | EastAfricans | 8.92E-04 | 7.13E-04 | 0.210637 |
| *BNC2* | rs10756819 | G | A | Oceanians | EastAfricans | 0.001476 | 6.84E-04 | **0.031069** |
| *BNC2* | rs12350739 | G | A | Oceanians | EastAfricans | 2.35E-04 | 9.97E-04 | 0.813429 |
| *DDB1* | rs11230664 | C | T | Oceanians | EastAfricans | 0.001111 | 6.56E-04 | 0.090412 |
| *DDB1* | rs7120594 | T | C | Oceanians | EastAfricans | 5.94E-04 | 6.45E-04 | 0.357396 |
| *DDB1* | rs7948623 | A | T | Oceanians | EastAfricans | -3.64E-04 | 6.43E-04 | 0.57195 |
| *DDB1* | rs1377457 | C | A | Oceanians | EastAfricans | 0.001174 | 6.59E-04 | 0.074815 |
| *TPCN2* | rs35264875 | A | T | Oceanians | EastAfricans | 4.90E-04 | 6.46E-04 | 0.448032 |
| *TPCN2* | rs3829241 | G | A | Oceanians | EastAfricans | 3.77E-04 | 7.10E-04 | 0.59576 |
| *TYR* | rs1042602 | C | A | Oceanians | EastAfricans | 2.35E-04 | 9.97E-04 | 0.813429 |
| *TYR* | rs1393350 | G | A | Oceanians | EastAfricans | 2.35E-04 | 9.97E-04 | 0.813429 |
| *TYR* | rs1126809 | G | A | Oceanians | EastAfricans | 2.35E-04 | 9.97E-04 | 0.813429 |
| *KITLG* | rs642742 | A | G | Oceanians | EastAfricans | 0.001297 | 6.52E-04 | **0.04667** |
| *KITLG* | rs12821256 | T | C | Oceanians | EastAfricans | -8.58E-04 | 8.47E-04 | 0.310831 |
| *DCT* | rs1407995 | T | C | Oceanians | EastAfricans | -5.38E-04 | 6.46E-04 | 0.404862 |
| *DCT* | rs2031526 | G | A | Oceanians | EastAfricans | 0.001103 | 6.63E-04 | 0.096146 |
| *SLC24A4* | rs12896399 | G | T | Oceanians | EastAfricans | 0.001184 | 6.56E-04 | 0.071209 |
| *SLC24A4* | rs2402130 | G | A | Oceanians | EastAfricans | 6.62E-04 | 6.49E-04 | 0.308068 |
| *OCA2* | rs2311843 | C | T | Oceanians | EastAfricans | 0.001748 | 7.13E-04 | **0.014191** |
| *OCA2* | rs1800414 | A | G | Oceanians | EastAfricans | 0.001612 | 8.42E-04 | 0.055664 |
| *OCA2* | rs1800404 | C | T | Oceanians | EastAfricans | 9.40E-04 | 6.66E-04 | 0.158276 |
| *OCA2* | rs1800401 | C | T | Oceanians | EastAfricans | -6.23E-04 | 8.55E-04 | 0.466227 |
| *HERC2* | rs1129038 | G | A | Oceanians | EastAfricans | 2.35E-04 | 9.97E-04 | 0.813429 |
| *HERC2* | rs12913832 | A | G | Oceanians | EastAfricans | 2.35E-04 | 9.97E-04 | 0.813429 |
| *HERC2* | rs916977 | A | G | Oceanians | EastAfricans | 0.001428 | 6.85E-04 | **0.036998** |
| *HERC2* | rs1667394 | G | A | Oceanians | EastAfricans | 0.001428 | 6.85E-04 | **0.036998** |
| *SLC24A5* | rs1426654 | G | A | Oceanians | EastAfricans | -9.40E-05 | 7.37E-04 | 0.898072 |
| *MYO5A* | rs4776053 | C | T | Oceanians | EastAfricans | -3.75E-04 | 7.23E-04 | 0.604181 |
| *MC1R* | rs2228479 | G | A | Oceanians | EastAfricans | 0.001295 | 8.48E-04 | 0.126915 |
| *MC1R* | rs885479 | G | A | Oceanians | EastAfricans | 8.67E-04 | 8.70E-04 | 0.319434 |
| *MFSD12* | rs56203814 | C | T | Oceanians | EastAfricans | -0.001207 | 8.41E-04 | 0.151188 |
| *MFSD12* | rs10424065 | C | T | Oceanians | EastAfricans | -0.002011 | 8.38E-04 | 0.016387 |
| *MFSD12* | rs6510760 | G | A | Oceanians | EastAfricans | -0.001219 | 6.65E-04 | 0.066782 |
| *MFSD12* | rs112332856 | T | C | Oceanians | EastAfricans | -0.001604 | 6.67E-04 | **0.016127** |
| *ASIP* | rs6119471 | G | C | Oceanians | EastAfricans | 0.00155 | 8.38E-04 | 0.06457 |
| *ASIP* | rs6058017 | G | A | Oceanians | EastAfricans | 0.001098 | 6.51E-04 | 0.091568 |
| *PIGU* | rs2378249 | A | G | Oceanians | EastAfricans | 3.67E-04 | 6.64E-04 | 0.580531 |
| *SLC45A2* | rs16891982 | C | G | Oceanians | WestAfricans | 9.61E-04 | 5.60E-04 | 0.086576 |
| *SLC45A2* | rs28777 | C | A | Oceanians | WestAfricans | 4.33E-04 | 5.24E-04 | 0.408818 |
| *SLC45A2* | rs26722 | C | T | Oceanians | WestAfricans | 5.63E-04 | 5.27E-04 | 0.286 |
| *EXOC2* | rs4959270 | C | A | Oceanians | WestAfricans | 9.84E-04 | 5.31E-04 | 0.063704 |
| *TYRP1* | rs1408799 | T | C | Oceanians | WestAfricans | -6.57E-04 | 5.76E-04 | 0.253421 |
| *TYRP1* | rs683 | A | C | Oceanians | WestAfricans | 4.44E-04 | 5.76E-04 | 0.441185 |
| *BNC2* | rs10756819 | G | A | Oceanians | WestAfricans | 8.03E-04 | 5.25E-04 | 0.125638 |
| *BNC2* | rs12350739 | G | A | Oceanians | WestAfricans | 3.40E-04 | 7.00E-04 | 0.626702 |
| *DDB1* | rs11230664 | C | T | Oceanians | WestAfricans | 2.80E-04 | 5.24E-04 | 0.592903 |
| *DDB1* | rs7120594 | T | C | Oceanians | WestAfricans | 1.52E-04 | 5.23E-04 | 0.771004 |
| *DDB1* | rs7948623 | A | T | Oceanians | WestAfricans | 2.70E-04 | 5.24E-04 | 0.605794 |
| *DDB1* | rs1377457 | C | A | Oceanians | WestAfricans | 2.99E-04 | 5.24E-04 | 0.568196 |
| *TPCN2* | rs35264875 | A | T | Oceanians | WestAfricans | 0.001402 | 5.36E-04 | **0.008884** |
| *TPCN2* | rs3829241 | G | A | Oceanians | WestAfricans | 2.27E-04 | 5.47E-04 | 0.677517 |
| *TYR* | rs1042602 | C | A | Oceanians | WestAfricans | 2.38E-04 | 6.92E-04 | 0.730883 |
| *TYR* | rs1393350 | G | A | Oceanians | WestAfricans | 8.29E-04 | 7.98E-04 | 0.299012 |
| *TYR* | rs1126809 | G | A | Oceanians | WestAfricans | 2.38E-04 | 6.92E-04 | 0.730883 |
| *KITLG* | rs642742 | A | G | Oceanians | WestAfricans | 0.001175 | 5.27E-04 | **0.025836** |
| *KITLG* | rs12821256 | T | C | Oceanians | WestAfricans | 8.29E-04 | 7.98E-04 | 0.299012 |
| *DCT* | rs1407995 | T | C | Oceanians | WestAfricans | -3.25E-04 | 5.24E-04 | 0.534328 |
| *DCT* | rs2031526 | G | A | Oceanians | WestAfricans | 7.87E-04 | 5.25E-04 | 0.133799 |
| *SLC24A4* | rs12896399 | G | T | Oceanians | WestAfricans | 0.001906 | 5.57E-04 | **6.26E-04** |
| *SLC24A4* | rs2402130 | G | A | Oceanians | WestAfricans | 6.51E-04 | 5.28E-04 | 0.217798 |
| *OCA2* | rs2311843 | C | T | Oceanians | WestAfricans | 0.001163 | 5.27E-04 | **0.027279** |
| *OCA2* | rs1800414 | A | G | Oceanians | WestAfricans | 0.001913 | 6.78E-04 | **0.004789** |
| *OCA2* | rs1800404 | C | T | Oceanians | WestAfricans | 3.90E-04 | 5.25E-04 | 0.457676 |
| *OCA2* | rs1800401 | C | T | Oceanians | WestAfricans | -9.50E-04 | 6.74E-04 | 0.158555 |
| *HERC2* | rs1129038 | G | A | Oceanians | WestAfricans | 1.29E-04 | 6.87E-04 | 0.850487 |
| *HERC2* | rs12913832 | A | G | Oceanians | WestAfricans | 1.29E-04 | 6.87E-04 | 0.850487 |
| *HERC2* | rs916977 | A | G | Oceanians | WestAfricans | 7.23E-04 | 5.24E-04 | 0.167892 |
| *HERC2* | rs1667394 | G | A | Oceanians | WestAfricans | 7.65E-04 | 5.25E-04 | 0.144519 |
| *SLC24A5* | rs1426654 | G | A | Oceanians | WestAfricans | -2.32E-04 | 5.77E-04 | 0.687656 |
| *MYO5A* | rs4776053 | C | T | Oceanians | WestAfricans | -3.85E-04 | 5.76E-04 | 0.504393 |
| *MC1R* | rs2228479 | G | A | Oceanians | WestAfricans | 0.001664 | 6.83E-04 | **0.014853** |
| *MC1R* | rs885479 | G | A | Oceanians | WestAfricans | 0.001326 | 7.00E-04 | 0.05814 |
| *MFSD12* | rs56203814 | C | T | Oceanians | WestAfricans | -0.001078 | 6.73E-04 | 0.109476 |
| *MFSD12* | rs10424065 | C | T | Oceanians | WestAfricans | -0.001234 | 6.73E-04 | 0.066848 |
| *MFSD12* | rs6510760 | G | A | Oceanians | WestAfricans | -5.80E-04 | 5.24E-04 | 0.268324 |
| *MFSD12* | rs112332856 | T | C | Oceanians | WestAfricans | -7.24E-04 | 5.25E-04 | 0.167624 |
| *ASIP* | rs6119471 | G | C | Oceanians | WestAfricans | 0.001799 | 6.73E-04 | **0.007547** |
| *ASIP* | rs6058017 | G | A | Oceanians | WestAfricans | 9.99E-04 | 5.29E-04 | 0.058665 |
| *PIGU* | rs2378249 | A | G | Oceanians | WestAfricans | -4.20E-05 | 5.30E-04 | 0.936817 |
| *SLC45A2* | rs16891982 | C | G | EastAsians | EastAfricans | -1.11E-04 | 7.02E-04 | 0.873917 |
| *SLC45A2* | rs28777 | C | A | EastAsians | EastAfricans | -3.32E-04 | 6.27E-04 | 0.596567 |
| *SLC45A2* | rs26722 | C | T | EastAsians | EastAfricans | 9.01E-04 | 6.39E-04 | 0.158605 |
| *EXOC2* | rs4959270 | C | A | EastAsians | EastAfricans | 8.07E-04 | 6.39E-04 | 0.206507 |
| *TYRP1* | rs1408799 | T | C | EastAsians | EastAfricans | -0.001057 | 6.29E-04 | 0.093125 |
| *TYRP1* | rs683 | A | C | EastAsians | EastAfricans | 0.001221 | 6.33E-04 | 0.053946 |
| *BNC2* | rs10756819 | G | A | EastAsians | EastAfricans | 0.001561 | 6.68E-04 | **0.019443** |
| *BNC2* | rs12350739 | G | A | EastAsians | EastAfricans | 1.24E-04 | 8.34E-04 | 0.882304 |
| *DDB1* | rs11230664 | C | T | EastAsians | EastAfricans | 0.002346 | 6.40E-04 | **2.48E-04** |
| *DDB1* | rs7120594 | T | C | EastAsians | EastAfricans | 0.001829 | 6.29E-04 | **0.003637** |
| *DDB1* | rs7948623 | A | T | EastAsians | EastAfricans | -0.002662 | 6.58E-04 | **5.20E-05** |
| *DDB1* | rs1377457 | C | A | EastAsians | EastAfricans | 0.003039 | 6.50E-04 | **3.00E-06** |
| *TPCN2* | rs35264875 | A | T | EastAsians | EastAfricans | -0.001222 | 6.35E-04 | 0.05446 |
| *TPCN2* | rs3829241 | G | A | EastAsians | EastAfricans | 9.51E-04 | 6.68E-04 | 0.154736 |
| *TYR* | rs1042602 | C | A | EastAsians | EastAfricans | -4.48E-04 | 8.61E-04 | 0.602762 |
| *TYR* | rs1393350 | G | A | EastAsians | EastAfricans | -3.19E-04 | 8.51E-04 | 0.708248 |
| *TYR* | rs1126809 | G | A | EastAsians | EastAfricans | -6.45E-04 | 8.84E-04 | 0.465599 |
| *KITLG* | rs642742 | A | G | EastAsians | EastAfricans | 0.001281 | 6.31E-04 | **0.0423** |
| *KITLG* | rs12821256 | T | C | EastAsians | EastAfricans | -0.001738 | 7.10E-04 | **0.014326** |
| *DCT* | rs1407995 | T | C | EastAsians | EastAfricans | -0.00104 | 6.28E-04 | 0.098027 |
| *DCT* | rs2031526 | G | A | EastAsians | EastAfricans | 0.001607 | 6.45E-04 | **0.012763** |
| *SLC24A4* | rs12896399 | G | T | EastAsians | EastAfricans | 7.97E-04 | 6.39E-04 | 0.212191 |
| *SLC24A4* | rs2402130 | G | A | EastAsians | EastAfricans | 0.001059 | 6.26E-04 | 0.090478 |
| *OCA2* | rs2311843 | C | T | EastAsians | EastAfricans | 0.002157 | 6.97E-04 | **0.001962** |
| *OCA2* | rs1800414 | A | G | EastAsians | EastAfricans | 0.002249 | 8.26E-04 | **0.006478** |
| *OCA2* | rs1800404 | C | T | EastAsians | EastAfricans | 0.001069 | 6.48E-04 | 0.099004 |
| *OCA2* | rs1800401 | C | T | EastAsians | EastAfricans | -6.41E-04 | 6.56E-04 | 0.327988 |
| *HERC2* | rs1129038 | G | A | EastAsians | EastAfricans | -2.22E-04 | 8.46E-04 | 0.79322 |
| *HERC2* | rs12913832 | A | G | EastAsians | EastAfricans | -3.19E-04 | 8.51E-04 | 0.708248 |
| *HERC2* | rs916977 | A | G | EastAsians | EastAfricans | 0.001082 | 6.68E-04 | 0.105386 |
| *HERC2* | rs1667394 | G | A | EastAsians | EastAfricans | 0.00108 | 6.68E-04 | 0.105724 |
| *SLC24A5* | rs1426654 | G | A | EastAsians | EastAfricans | -3.49E-04 | 6.60E-04 | 0.597169 |
| *MYO5A* | rs4776053 | C | T | EastAsians | EastAfricans | 4.53E-04 | 6.39E-04 | 0.478246 |
| *MC1R* | rs2228479 | G | A | EastAsians | EastAfricans | 0.001758 | 8.26E-04 | **0.033387** |
| *MC1R* | rs885479 | G | A | EastAsians | EastAfricans | 0.002282 | 8.26E-04 | **0.005752** |
| *MFSD12* | rs56203814 | C | T | EastAsians | EastAfricans | -0.00251 | 8.31E-04 | **0.002534** |
| *MFSD12* | rs10424065 | C | T | EastAsians | EastAfricans | -0.002891 | 6.99E-04 | **3.60E-05** |
| *MFSD12* | rs6510760 | G | A | EastAsians | EastAfricans | -0.002354 | 6.49E-04 | **2.86E-04** |
| *MFSD12* | rs112332856 | T | C | EastAsians | EastAfricans | -0.003299 | 6.60E-04 | **1.00E-06** |
| *ASIP* | rs6119471 | G | C | EastAsians | EastAfricans | 0.002852 | 8.29E-04 | **5.79E-04** |
| *ASIP* | rs6058017 | G | A | EastAsians | EastAfricans | 9.13E-04 | 6.27E-04 | 0.144925 |
| *PIGU* | rs2378249 | A | G | EastAsians | EastAfricans | 4.46E-04 | 6.39E-04 | 0.485309 |
| *SLC45A2* | rs16891982 | C | G | EastAsians | WestAfricans | 3.00E-04 | 5.36E-04 | 0.575049 |
| *SLC45A2* | rs28777 | C | A | EastAsians | WestAfricans | -9.20E-05 | 5.07E-04 | 0.85544 |
| *SLC45A2* | rs26722 | C | T | EastAsians | WestAfricans | 7.93E-04 | 5.08E-04 | 0.118286 |
| *EXOC2* | rs4959270 | C | A | EastAsians | WestAfricans | 1.93E-04 | 5.06E-04 | 0.703733 |
| *TYRP1* | rs1408799 | T | C | EastAsians | WestAfricans | -7.11E-04 | 5.09E-04 | 0.16208 |
| *TYRP1* | rs683 | A | C | EastAsians | WestAfricans | 7.02E-04 | 5.11E-04 | 0.169651 |
| *BNC2* | rs10756819 | G | A | EastAsians | WestAfricans | 8.70E-04 | 5.07E-04 | 0.086276 |
| *BNC2* | rs12350739 | G | A | EastAsians | WestAfricans | 2.52E-04 | 5.48E-04 | 0.64552 |
| *DDB1* | rs11230664 | C | T | EastAsians | WestAfricans | 0.001253 | 5.07E-04 | **0.013528** |
| *DDB1* | rs7120594 | T | C | EastAsians | WestAfricans | 0.001126 | 5.07E-04 | **0.026524** |
| *DDB1* | rs7948623 | A | T | EastAsians | WestAfricans | -0.001541 | 5.31E-04 | **0.003738** |
| *DDB1* | rs1377457 | C | A | EastAsians | WestAfricans | 0.001768 | 5.13E-04 | **5.67E-04** |
| *TPCN2* | rs35264875 | A | T | EastAsians | WestAfricans | 5.30E-05 | 5.24E-04 | 0.919283 |
| *TPCN2* | rs3829241 | G | A | EastAsians | WestAfricans | 6.80E-04 | 5.09E-04 | 0.182182 |
| *TYR* | rs1042602 | C | A | EastAsians | WestAfricans | -3.00E-04 | 5.65E-04 | 0.59469 |
| *TYR* | rs1393350 | G | A | EastAsians | WestAfricans | 3.92E-04 | 6.82E-04 | 0.565332 |
| *TYR* | rs1126809 | G | A | EastAsians | WestAfricans | -4.55E-04 | 5.86E-04 | 0.437031 |
| *KITLG* | rs642742 | A | G | EastAsians | WestAfricans | 0.001162 | 5.07E-04 | **0.021928** |
| *KITLG* | rs12821256 | T | C | EastAsians | WestAfricans | 1.35E-04 | 7.08E-04 | 0.8484 |
| *DCT* | rs1407995 | T | C | EastAsians | WestAfricans | -7.21E-04 | 5.06E-04 | 0.154743 |
| *DCT* | rs2031526 | G | A | EastAsians | WestAfricans | 0.001185 | 5.08E-04 | **0.019612** |
| *SLC24A4* | rs12896399 | G | T | EastAsians | WestAfricans | 0.001601 | 5.41E-04 | **0.00308** |
| *SLC24A4* | rs2402130 | G | A | EastAsians | WestAfricans | 9.65E-04 | 5.07E-04 | 0.057024 |
| *OCA2* | rs2311843 | C | T | EastAsians | WestAfricans | 0.001485 | 5.10E-04 | **0.00356** |
| *OCA2* | rs1800414 | A | G | EastAsians | WestAfricans | 0.002416 | 6.63E-04 | **2.69E-04** |
| *OCA2* | rs1800404 | C | T | EastAsians | WestAfricans | 4.91E-04 | 5.07E-04 | 0.332155 |
| *OCA2* | rs1800401 | C | T | EastAsians | WestAfricans | -9.64E-04 | 5.13E-04 | 0.06006 |
| *HERC2* | rs1129038 | G | A | EastAsians | WestAfricans | -2.31E-04 | 5.43E-04 | 0.670853 |
| *HERC2* | rs12913832 | A | G | EastAsians | WestAfricans | -3.07E-04 | 5.48E-04 | 0.575425 |
| *HERC2* | rs916977 | A | G | EastAsians | WestAfricans | 4.50E-04 | 5.07E-04 | 0.374526 |
| *HERC2* | rs1667394 | G | A | EastAsians | WestAfricans | 4.92E-04 | 5.07E-04 | 0.332331 |
| *SLC24A5* | rs1426654 | G | A | EastAsians | WestAfricans | -4.32E-04 | 5.12E-04 | 0.398198 |
| *MYO5A* | rs4776053 | C | T | EastAsians | WestAfricans | 2.68E-04 | 5.07E-04 | 0.597823 |
| *MC1R* | rs2228479 | G | A | EastAsians | WestAfricans | 0.002028 | 6.63E-04 | **0.00222** |
| *MC1R* | rs885479 | G | A | EastAsians | WestAfricans | 0.002441 | 6.63E-04 | **2.32E-04** |
| *MFSD12* | rs56203814 | C | T | EastAsians | WestAfricans | -0.002104 | 6.63E-04 | **0.001509** |
| *MFSD12* | rs10424065 | C | T | EastAsians | WestAfricans | -0.001928 | 5.63E-04 | **6.22E-04** |
| *MFSD12* | rs6510760 | G | A | EastAsians | WestAfricans | -0.001474 | 5.07E-04 | **0.003679** |
| *MFSD12* | rs112332856 | T | C | EastAsians | WestAfricans | -0.00206 | 5.16E-04 | **6.60E-05** |
| *ASIP* | rs6119471 | G | C | EastAsians | WestAfricans | 0.002826 | 6.63E-04 | **2.00E-05** |
| *ASIP* | rs6058017 | G | A | EastAsians | WestAfricans | 8.54E-04 | 5.07E-04 | 0.091702 |
| *PIGU* | rs2378249 | A | G | EastAsians | WestAfricans | 2.00E-05 | 5.07E-04 | 0.967942 |
| *SLC45A2* | rs16891982 | C | G | EastAfricans | WestAfricans | 3.88E-04 | 3.41E-04 | 0.255086 |
| *SLC45A2* | rs28777 | C | A | EastAfricans | WestAfricans | 1.69E-04 | 1.81E-04 | 0.351565 |
| *SLC45A2* | rs26722 | C | T | EastAfricans | WestAfricans | 8.40E-05 | 2.10E-04 | 0.689776 |
| *EXOC2* | rs4959270 | C | A | EastAfricans | WestAfricans | -4.43E-04 | 2.06E-04 | **0.03162** |
| *TYRP1* | rs1408799 | T | C | EastAfricans | WestAfricans | 1.22E-04 | 1.82E-04 | 0.503424 |
| *TYRP1* | rs683 | A | C | EastAfricans | WestAfricans | -2.59E-04 | 1.85E-04 | 0.160278 |
| *BNC2* | rs10756819 | G | A | EastAfricans | WestAfricans | -3.59E-04 | 2.59E-04 | 0.165122 |
| *BNC2* | rs12350739 | G | A | EastAfricans | WestAfricans | 1.55E-04 | 4.99E-04 | 0.756223 |
| *DDB1* | rs11230664 | C | T | EastAfricans | WestAfricans | -5.95E-04 | 2.06E-04 | **0.003833** |
| *DDB1* | rs7120594 | T | C | EastAfricans | WestAfricans | -3.15E-04 | 1.83E-04 | 0.084296 |
| *DDB1* | rs7948623 | A | T | EastAfricans | WestAfricans | 5.57E-04 | 1.79E-04 | **0.001852** |
| *DDB1* | rs1377457 | C | A | EastAfricans | WestAfricans | -6.26E-04 | 2.11E-04 | **0.003034** |
| *TPCN2* | rs35264875 | A | T | EastAfricans | WestAfricans | 0.001016 | 2.16E-04 | **3.00E-06** |
| *TPCN2* | rs3829241 | G | A | EastAfricans | WestAfricans | -6.90E-05 | 2.63E-04 | 0.792385 |
| *TYR* | rs1042602 | C | A | EastAfricans | WestAfricans | 5.30E-05 | 4.89E-04 | 0.914067 |
| *TYR* | rs1393350 | G | A | EastAfricans | WestAfricans | 6.43E-04 | 6.29E-04 | 0.306763 |
| *TYR* | rs1126809 | G | A | EastAfricans | WestAfricans | 5.30E-05 | 4.89E-04 | 0.914067 |
| *KITLG* | rs642742 | A | G | EastAfricans | WestAfricans | 1.53E-04 | 1.92E-04 | 0.423979 |
| *KITLG* | rs12821256 | T | C | EastAfricans | WestAfricans | 0.001505 | 4.73E-04 | **0.001479** |
| *DCT* | rs1407995 | T | C | EastAfricans | WestAfricans | 9.90E-05 | 1.85E-04 | 0.594727 |
| *DCT* | rs2031526 | G | A | EastAfricans | WestAfricans | -8.20E-05 | 2.21E-04 | 0.712095 |
| *SLC24A4* | rs12896399 | G | T | EastAfricans | WestAfricans | 9.73E-04 | 2.81E-04 | **5.26E-04** |
| *SLC24A4* | rs2402130 | G | A | EastAfricans | WestAfricans | 1.30E-04 | 1.78E-04 | 0.464636 |
| *OCA2* | rs2311843 | C | T | EastAfricans | WestAfricans | -2.14E-04 | 3.06E-04 | 0.483851 |
| *OCA2* | rs1800414 | A | G | EastAfricans | WestAfricans | 6.43E-04 | 6.29E-04 | 0.306763 |
| *OCA2* | rs1800404 | C | T | EastAfricans | WestAfricans | -3.51E-04 | 2.24E-04 | 0.11748 |
| *OCA2* | rs1800401 | C | T | EastAfricans | WestAfricans | -4.59E-04 | 2.24E-04 | 0.040104 |
| *HERC2* | rs1129038 | G | A | EastAfricans | WestAfricans | -5.60E-05 | 4.81E-04 | 0.90728 |
| *HERC2* | rs12913832 | A | G | EastAfricans | WestAfricans | -5.60E-05 | 4.81E-04 | 0.90728 |
| *HERC2* | rs916977 | A | G | EastAfricans | WestAfricans | -4.02E-04 | 2.59E-04 | 0.120361 |
| *HERC2* | rs1667394 | G | A | EastAfricans | WestAfricans | -3.59E-04 | 2.59E-04 | 0.165122 |
| *SLC24A5* | rs1426654 | G | A | EastAfricans | WestAfricans | -1.57E-04 | 2.39E-04 | 0.509837 |
| *MYO5A* | rs4776053 | C | T | EastAfricans | WestAfricans | -8.90E-05 | 2.08E-04 | 0.66729 |
| *MC1R* | rs2228479 | G | A | EastAfricans | WestAfricans | 6.43E-04 | 6.29E-04 | 0.306763 |
| *MC1R* | rs885479 | G | A | EastAfricans | WestAfricans | 6.43E-04 | 6.29E-04 | 0.306763 |
| *MFSD12* | rs56203814 | C | T | EastAfricans | WestAfricans | -1.27E-04 | 1.86E-04 | 0.494038 |
| *MFSD12* | rs10424065 | C | T | EastAfricans | WestAfricans | 3.51E-04 | 1.78E-04 | **0.048901** |
| *MFSD12* | rs6510760 | G | A | EastAfricans | WestAfricans | 3.81E-04 | 2.24E-04 | 0.089162 |
| *MFSD12* | rs112332856 | T | C | EastAfricans | WestAfricans | 5.39E-04 | 2.23E-04 | **0.015703** |
| *ASIP* | rs6119471 | G | C | EastAfricans | WestAfricans | 5.78E-04 | 1.79E-04 | **0.001269** |
| *ASIP* | rs6058017 | G | A | EastAfricans | WestAfricans | 1.35E-04 | 1.82E-04 | 0.458646 |
| *PIGU* | rs2378249 | A | G | EastAfricans | WestAfricans | -3.31E-04 | 2.07E-04 | 0.109172 |

### Text S1. The effects of migration and substructure

In this section, we examined how the estimated selection difference 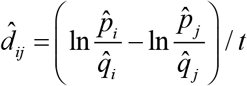 is affected by population migration and substructure in theory. Here, 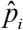 and 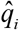 are the observed derived and ancestral allele frequencies in the population *i*, respectively; 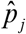. and 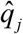 are the observed derived and ancestral allele frequencies in the population *j*, respectively; and *t* is the divergence time from populations *i* and *j* to their most recent common ancestor. We demonstrated that 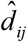 provides a lower bound of selection difference between populations *i* and *j* when migration or substructure exists. We first provide two inequalities that will be used later.

#### Inequality 1

If *a* > *b* > 0, *c* > *d* > 0, then *ac* > *bd* and 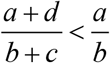.

Proof: *a* > *b* > 0, *c* > *d* > 0, then *ac* > *bc*, *bc* > *bd*. Therefore, *ac* > *bd*. Furthermore, *ac* + *ab* > *bd* + *ab*, which is the same as *a*(*b* + *c*)>*b*(*a* + *d*). Therefore, 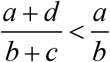.

#### Inequality 2

If *a*_1_ > 0, *a*_2_ > 0, L, *a_n_* > 0, 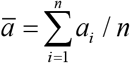, then max {*a*_1_, *a*_2_, L, *a_n_*} ≥ *ā*, min{*a*_1_, *a*_2_, L, *a_n_*} ≤ *ā*.

Proof: Let *a*_max_ = max{*a*_1_, *a*_2_,L, *a_n_*}, then

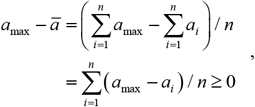

because *a*_max_ ≥ *a_i_*.

Let *a*_min_ = min {*a*_1_,*a*_2_,L, *a_n_*}, then

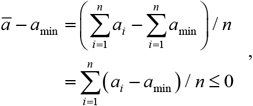

because *a_min_* ≤ *a_i_*.

For the proofs in below, we assume 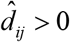 without loss of generality. If 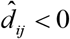, then we can exchange *i* and *j*, and still obtain 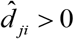.

#### The effect of migration

Suppose there is *α* proportion of individuals in the population *i*, which actually come from the population *j*; also, there is *β* proportion of individuals in the population *j*, which actually come from the population *i*. Here, 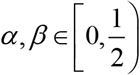, because we assume migrants should not become the majority of another population. Then we have

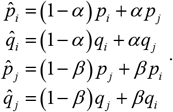

Here, *p_i_* and *q_i_* are the true derived and ancestral allele frequencies in the population *i*, respectively; *p_j_* and *q_j_* are the true derived and ancestral allele frequencies in the population *j*, respectively. Therefore,

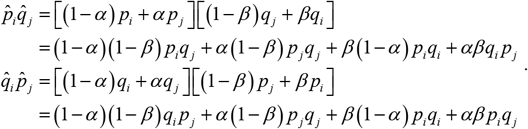

Further, we have

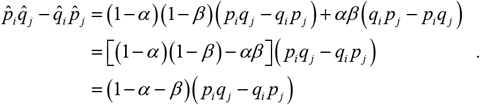

Because 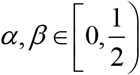, then 1 − *α* − *β* > 0; and 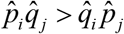, therefore, *p_i_q_j_* − *q_i_p_j_* > 0.

We also have 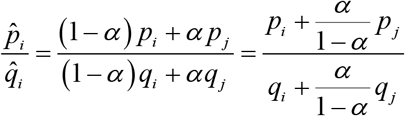. Because *p_i_q_j_* − *q_i_p_j_* > 0, we have 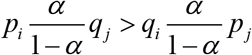. From Inequality 1, we know 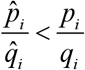. Similarly, we also have 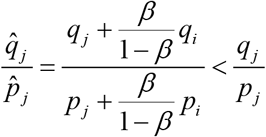, therefore, 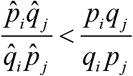. According to the monotony of the logarithmic function, we have 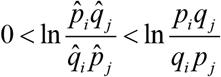; thus, 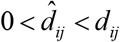. In other words, if migration exists between populations *i* and *j*, the estimated selection difference is lower than the true value.

#### The effect of substructure

**Scenario 1:** The population *j* has *k* subpopulations.

If the population *j* has *k* subpopulations, then 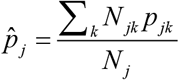. Here, *p_jk_* is the derived allele frequency in the subpopulation *k* of the population *j*. And *N_j_* is the population size of the population *j*, 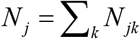. We denote the minimum of *p_jk_* as min (*p_jk_*). Because *q_jk_* =1 − *p_jk_*, then max (*q_jk_*) = 1 − min (*p_jk_*). Based on Inequality 2, 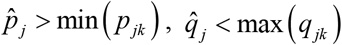. Therefore, 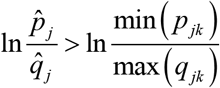. We have 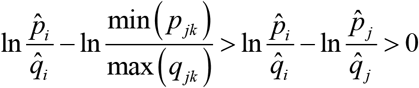.

**Scenario 2:** The population *i* has *l* subpopulations.

If the population *i* has *l* subpopulations, then 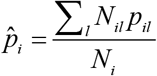. We denote the maximum of *p_il_* as max (*p_il_*). Then 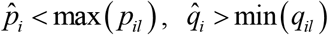. Therefore, 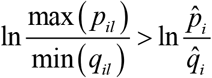, and we have 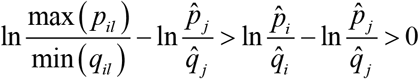.

**Scenario 3:** The population *i* has *l* subpopulations, and the population *j* has *k* subpopulations Based on Scenario 1 and 2, we have 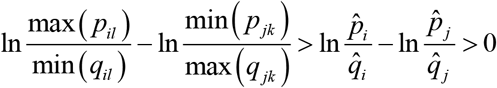. In summary, if populations *i* and *j* have subpopulations, and their estimated selection difference is larger than zero, then at least one pair of their subpopulations has selection difference larger than zero. Moreover, the estimated difference is smaller than the largest difference between subpopulations.

### Text S2. A sample script for simulation with SLiM 2

~~~
initialize() {
      initializeMutationRate(1e-8);
      initializeMutationType(“m1”, 0.5, “f”, 0.0);
      initializeMutationType(“m2”, 0.5, “f”, 0.0);
      m1.mutationStackPolicy = “f”;
      m2.mutationStackPolicy = “f”;
      initializeGenomicElementType(“g1”, m1, 1.0);
      initializeGenomicElement(g1, 0, 999999);
      initializeRecombinationRate(1e-8); }
1 { sim.addSubpop(“p1”, 12000); }
1 late() {
      // Set the initial frequency of the beneficial allele to 0.001
      targets = sample(p1.genomes, 24);
      targets.addNewDrawnMutation(m2, 499999); }
300 {
      // Add West Africans
      sim.addSubpopSplit(“p2”, 12000, p1); }
1000 {
      // Add East Africans
      sim.addSubpopSplit(“p3”, 12000, p1);
      p1.setSubpopulationSize(5000); }
1600 {
      // Add Oceanians
      sim.addSubpopSplit(“p6”, 5000, p1); }
2000 {
      // Add Europeans
      sim.addSubpopSplit(“p4”, 5000, p1); }
2300 {
      // Add North Asians and p1 becomes East Asians
      sim.addSubpopSplit(“p5”, 5000, p1); }
3300:3600 {
      t = sim.generation − 3300;
      // Expansion in Europeans and North Asians
      // Final effective population size becomes ~10,000
      p4_size = round(5000 * exp(0.0023 * t));
      // Expansion in Asians
      // Final effective population size becomes ~20,000
      p1_size = round(5000 * exp(0.0046 * t));
      p4.setSubpopulationSize(asInteger(p4_size));
      p5.setSubpopulationSize(asInteger(p4_size));
      p1.setSubpopulationSize(asInteger(p1_size)); }
// s1 = −0.0188/31*2 = −0.001213
301:3600 fitness(m2, p2) { return 0.998787; }
// s10 = 0.00398/31*2 = 0.0002568
301:1000 fitness(m2, p1) { return 1.000257; }
// s2 = −0.0204/31*2 = −0.001316
1001:3600 fitness(m2, p3) { return 0.998684; }
// s9 = 0.00813/31*2 = 0.0005245
1001:1600 fitness(m2, p1) { return 1.000525; }
// s3 = −0.00614/31*2 = −0.0003961
1601:3600 fitness(m2, p6) { return 0.999603; }
// s8 = 0.00665/31*2 = 0.0004290
1601:2100 fitness(m2, p1) { return 1.000429; }
// s4 = 0.0249/31*2 = 0.001606
2001:3600 fitness(m2, p4) { return 1.001613; }
// s7 = 0.000218/31*2 = 0.0000141
2001:2300 fitness(m2, p1) { return 1.000014; }
// s5 = 0.00622/31*2 = 0.0004013
2301:3600 fitness(m2, p5) { return 1.000401; }
// s6 = −0.00480/31*2 = −0.0003097
2301:3600 fitness(m2, p1) { return 0.999690; }
3600 late() {
    mut1 = sim.mutationsOfType(m2);
    cat(“Count:”
      + “\t” + “WAF\t” + sim.mutationCounts(p2, mut1) + “\t”
      + “\t” + “EAF\t” + sim.mutationCounts(p3, mut1) + “\t”
      + “\t” + “OCN\t” + sim.mutationCounts(p6, mut1) + “\t”
      + “\t” + “EUR\t” + sim.mutationCounts(p4, mut1) + “\t”
      + “\t” + “NAS\t” + sim.mutationCounts(p5, mut1) + “\t”
      + “\t”+ “EAS\t” + sim.mutationCounts(p1, mut1)
      + “\n”);
sim.simulationFinished(); }
~~~

